# Transposable element variation inferred from long-read sequences in wild house mice from temperate and tropical environments

**DOI:** 10.64898/2026.08.21.746367

**Authors:** Yocelyn T. Gutiérrez-Guerrero, Athmaja Viswanath, Simon Orozco-Arias, Marta Coronado-Zamora, Jingtao Lilue, Josefa González, Michael W. Nachman

**Author notes:** Corresponding authors: Yocelyn T. Gutiérrez-Guerrero and Michael W. Nachman, 3101 Valley Life Sciences Building, Museum of Vertebrate Zoology, University of California, Berkeley, Berkeley, CA, 94720; Phone for Michael Nachman.

## Abstract

Transposable elements (TEs) constitute a large fraction of mammalian genomes yet their contribution to variation among individuals within natural populations remains largely unexplored. While most TE insertions are deleterious, some may be beneficial and contribute to adaptation. We characterized TE variation and assessed its potential adaptive role using long-read whole-genome sequencing of wild-caught house mice (*Mus musculus domesticus*) sampled from two populations inhabiting contrasting temperate and tropical environments and differing in morphology, physiology, and behavior. We sequenced 10 mice from each population and created highly contiguous de-novo genome assemblies for each individual, allowing us to identify TEs that are not present in the mouse reference genome and to characterize individual variation. By performing manual TE curation, we identified 506 non-redundant TE consensus sequences among all mice. On average, each wild mouse genome contained 1.47 million TE insertions, ∼4% of which were polymorphic among individuals. A small fraction of these polymorphic TE insertions were present in high frequency in just one of the populations, consistent with positive natural selection. Using liver RNA-seq in natural populations and in laboratory crosses, we studied gene expression at genes adjacent to polymorphic TEs. This identified a small set of TEs that are associated with the expression of nearby genes in a population-specific manner, nearly all of which showed independent signatures of positive selection. Together, these results provide the first detailed assessment of TE variation in natural populations of house mice and identify a small set of TE insertions that likely contribute to environmental adaptation.

## Introduction

Transposable elements (TEs) make up a very large fraction of mammalian genomes, accounting for 30% - 60% of the total genome size in most species (Osmanski et al. 2023). Despite their large contribution to total genome size, the role of TEs in genetic and phenotypic variation among individuals within populations is still largely unknown. TEs are selfish genetic elements that can be mobilized in the germline and thereby increase in frequency in a population (Bourque et al. 2018). TE insertions that have become fixed among all individuals in a population do not contribute to inter-individual variation. However, TE insertions can also be present in some individuals and not in others, and the extent of this variation in TE content in natural populations has not been well described.

Difficulty in accurately characterizing TE content stems in part from the challenges of mapping elements that are present in high numbers of copies. This problem is particularly acute in mammals. For example, the human genome contains over 1 million *Alu* sequences (Batzer and Deininger 2002). Another challenge comes from mapping variants to a single reference genome. Complex repetitive sequences will often be misplaced or unmapped when the sampled genome differs substantially from the reference genome (Yang et al. 2019). The availability of long-read sequencing technology helps ameliorate these problems and has been recently used to accurately characterize TE variation mostly in plants (Shahid and Slotkin 2020; Quadrana and Herdenson 2025) but also in natural populations of flies (Rech et al. 2022) and mosquitoes (Vargas-Chavez et al. 2022) and among laboratory strains of mice (Ferraj et al. 2023).

Most TE insertions are expected to be neutral or deleterious and therefore at low frequencies in populations (Charlesworth and Langley 1989). However, several striking examples of adaptation have been tied to the insertion of a transposable element that has risen to high frequencies in particular populations. For example, Daborn et al. (2002) showed that resistance to insecticide in *Drosophila melanogaster* is due to the insertion of an *Accord* TE upstream of the cytochrome P450 gene *Cyp6g1* leading to increased expression. In *D. simulans*, a *Doc* TE has inserted upstream of the same *Cyp6g1* gene and is associated with signatures of positive selection and increased expression, although the association with insecticide resistance is less clear (Schlenke and Begun 2004; Brookfield 2004). The *carbonaria* mutation in peppered moths that is associated with industrial melanism is due to the insertion of a TE in an intron of the *cortex* gene (van’t Hof et al. 2016). Polymorphic TE insertions have also been associated with increased resistance to oxidative stress and to bacterial infection in *D. melanogaster* (Guio et al. 2018; Ullastres et al. 2021; Merenciano and González 2023). These individual examples demonstrate that TE insertions may sometimes be adaptive (Casacuberta and González 2013; Hayward and Gilbert 2022).

Motivated by the recognition that some TE insertions may be beneficial, a number of studies have conducted genome-wide screens for TE insertions that are at high frequencies or associated with other signatures of positive selection (e.g. Gonzalez et al. 2008; Quadrana et al. 2016; Zhang and Tautz et al. 2021; Rech et al. 2019, 2022). TE insertions have the potential to modify gene expression by upregulating or downregulating nearby genes through several mechanisms including the insertion of novel *cis*-regulatory elements, the disruption of reading frames, the creation of new isoforms, and the modification of promoters (Feschotte 2008; Ellison and Bachtrog 2012; Coronado-Zamora and González, 2023; Galbraith and Hayward 2023; Feschotte 2026). Thus some studies of TE variation have sought to connect individual insertions to the expression of nearby genes (e.g. Gonzalez et al. 2008; Quadrana et al. 2016; Rech et al. 2022; Raingeval et al. 2024). For example, in *Arabidopsis* LTR retrotransposons have been shown to regulate the flowering C locus under heat stress (Raingeval et al. 2024). In *D. melanogaster* populations from five different climatic regions, TE polymorphisms are associated with the expression of nearby genes, such as *Cyp6a17*, which is involved in thermoregulation and may contribute to adaptation to varying environmental conditions (Rech et al. 2022).

The Western house mouse (*Mus musculus domesticus*) provides an excellent mammalian model for investigating the potential role of TEs in adaptation. Distributed globally in association with humans, house mice have successfully colonized a wide range of habitats, evolving morphological, physiological and behavioral phenotypes in response to diverse environmental pressures (Lynch 1992; Boursot et al. 1993; Phifer-Rixey et al. 2018). Studies in natural populations across the Americas have documented adaptive evolution along latitudinal gradients and contrasting thermal environments (Lynch 1992; Phifer-Rixey et al. 2018; Ferris et al. 2021; Gutiérrez-Guerrero et al. 2024). For example, mice from more northern and colder populations tend to have larger bodies and shorter extremities compared to mice from equatorial regions, consistent with Bergmann’s and Allen’s rules (Ballinger and Nachman 2022). Expression studies in F1 hybrids generated from crosses between wild-derived strains adapted to warm and cold climates have shown that adaptive gene expression divergence is driven mainly by *cis*-regulatory changes (Ballinger et al. 2023; Durkin et al. 2024). Despite compelling evidence that regulatory variation contributes to environmental adaptation in house mice and despite the potential for TE insertions to alter gene expression, the role of TE polymorphism in shaping gene regulation and adaptation in natural populations of house mice remains unexplored, primarily due to the repetitive architecture of TEs and the computational challenges associated with their analysis in complex genomes.

To circumvent these challenges, we sequenced the whole genomes of 10 wild mice from a temperate population (New Hampshire and Vermont, USA) and 10 wild mice from a tropical population (Manaus, Brazil) using HiFi PacBio long-read sequencing. Previous studies of TE variation in house mice described differences among laboratory strains using short-read (Nellåker et al. 2012) or long-read sequences (Ferraj et al. 2023). These studies identified TEs using the Dfam database which may not contain all naturally-occurring TE diversity. In contrast, we generated high quality genome assemblies for wild-caught mice and constructed a *de novo* manually curated library of consensus TE sequences for comprehensive TE annotation. Using this new genomic resource, we characterized the TE landscape across individual genomes and quantified the variability observed in natural populations. This allowed us to identify novel TEs, estimate the TE copy frequencies in the two populations, and identify both shared and unique TE polymorphisms, including high-frequency TE insertions that were specific to each population. We then generated RNA-seq data from the 20 wild-caught mice to study how TE insertion polymorphisms influence gene expression in natural populations, and we identified *cis*-regulatory expression differences of adjacent genes by studying allele-specific expression patterns in F1 hybrids of crosses between lab-reared tropical and temperate mice. These analyses identified a small set of insertions that likely contribute to environmental adaptation in tropical and temperate house mice.

## Results

### Long-read genome assemblies for wild-caught mice and wild-derived inbred strains

We generated long-read whole-genome sequences for 20 wild-caught mice: 10 from Manaus, Brazil and 10 from a locality at the border of New Hampshire and Vermont (hereafter referred to as NH-VT) (Table 1; Fig. 1a-b; Supplementary Table 1-2). The mean sequencing coverage was 28x, with consistently high read quality (mean Q34, Supplementary Table 2). To further characterize genomic variation associated with tropical and temperate environments, and to connect that variation with gene expression in laboratory studies (see below), we included additional long-read whole genome sequences from two wild-derived inbred strains of mice: SARA and MANA. SARA is a fully inbred strain derived from wild mice caught in Saratoga Springs, NY, a location geographically close to and genetically similar to the NH-VT population (Durkin et al. 2024). The SARA whole-genome sequence was taken from Dumont et al. (2024) and consists of PacBio reads at 10x coverage. The MANA strain is fully inbred and derives from wild-caught mice from the same Manaus population analyzed here. We sequenced this strain using PacBio reads at 41x coverage. Sequencing details and genome assembly statistics for all 22 individuals are given in Supplementary Tables 2 and 3.

**Table 1.**
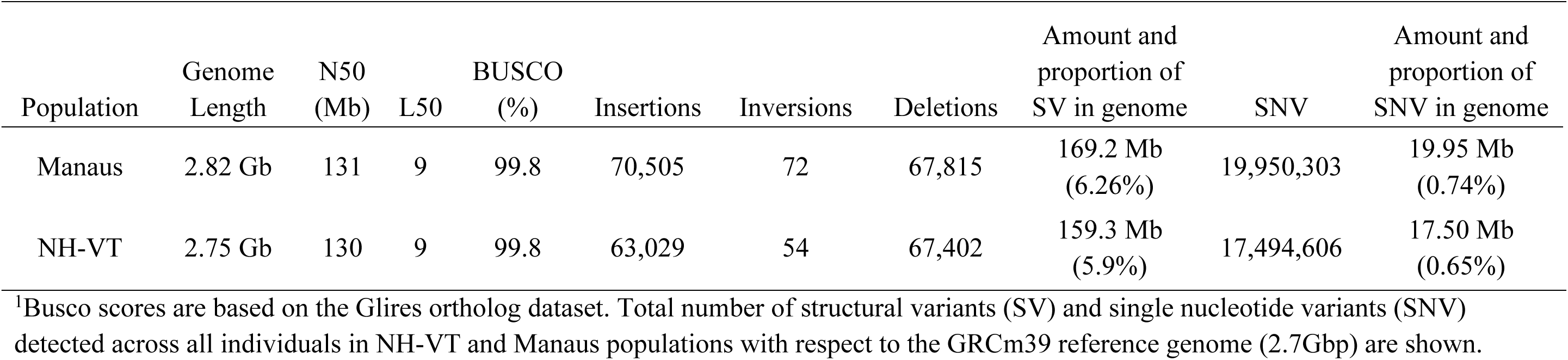
Whole genome assembly metrics and summaries of variation^1^.

| Population | Genome Length | N50 (Mb) | L50 | BUSCO (%) | Insertions | Inversions | Deletions | Amount and proportion of SV in genome | SNV | Amount and proportion of SNV in genome |
| --- | --- | --- | --- | --- | --- | --- | --- | --- | --- | --- |
| Manaus | 2.82 Gb | 131 | 9 | 99.8 | 70,505 | 72 | 67,815 | 169.2 Mb (6.26%) | 19,950,303 | 19.95 Mb (0.74%) |
| NH-VT | 2.75 Gb | 130 | 9 | 99.8 | 63,029 | 54 | 67,402 | 159.3 Mb (5.9%) | 17,494,606 | 17.50 Mb (0.65%) |

<sup>1</sup>Busco scores are based on the Glires ortholog dataset. Total number of structural variants (SV) and single nucleotide variants (SNV) detected across all individuals in NH-VT and Manaus populations with respect to the GRCm39 reference genome (2.7Gbp) are shown.

**Figure 1.**
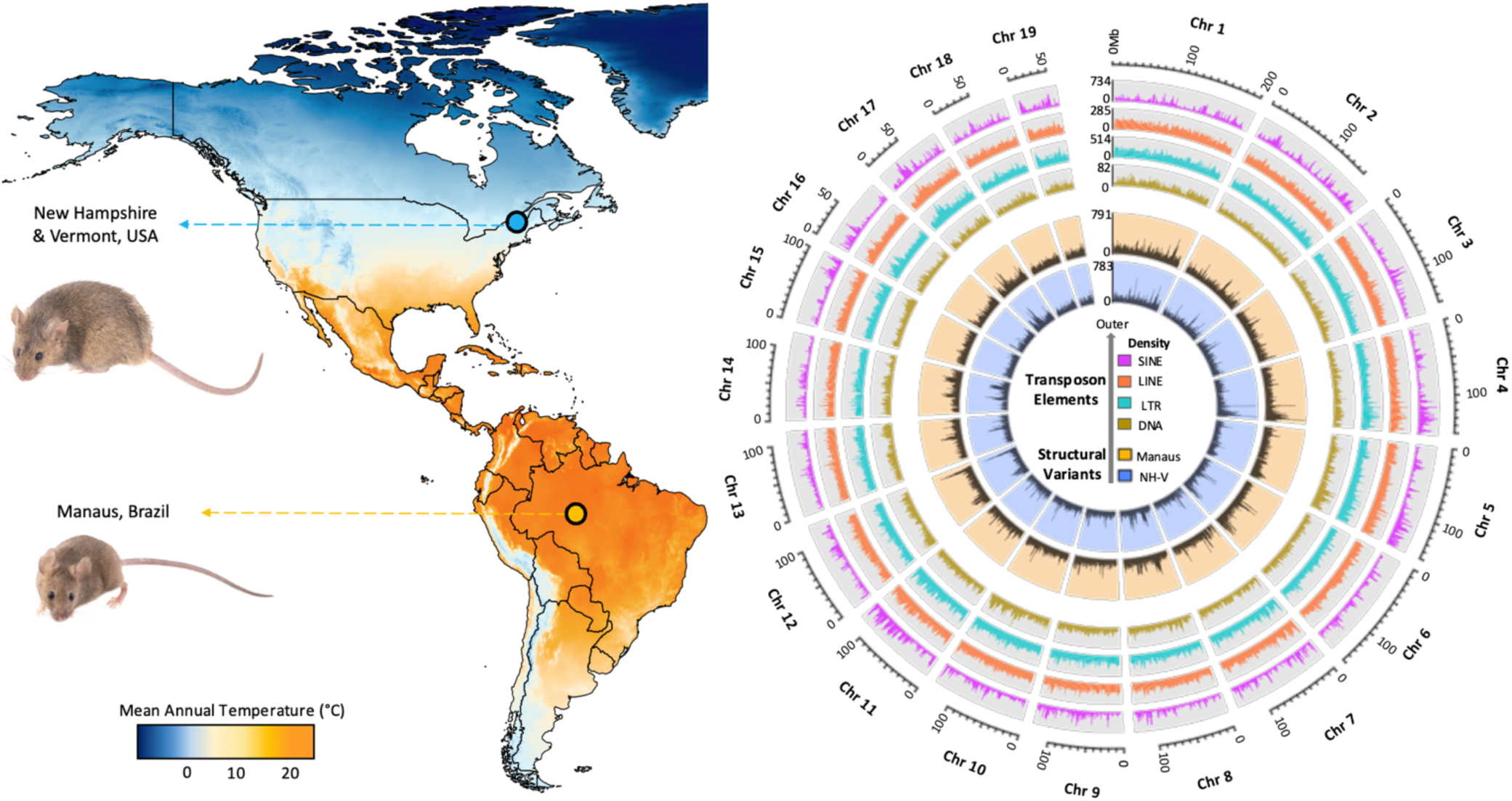
Sampling localities and summary of TE variation in wild mice. a) Map showing mean annual temperature of the temperate and tropical populations sampled: New Hampshire and Vermont, USA (NH-VT), and Manaus, Brazil. The wild-derived inbred strain SARA from a temperate population is larger and has shorter ears, tails, and limbs compared to the wild-derived inbred strain MANA from a tropical population, reflecting thermoregulatory adaptations to different climates (photo credit: Prakrit Jain). b) Average number of TEs annotated and structural variants discovered across all individuals sampled.

We constructed *de novo* whole-genome assemblies for each individual using hifiasm (Cheng et al. 2021). Scaffold N50 values ranged from 128 to 136 Mb, and L50 values ranged from 9 to 10. BUSCO analysis using the Glires ortholog dataset revealed an assembly with 99.8% complete single-copy orthologs (Table 1; Supplementary Table 3) indicating that these genomes are highly complete. For each individual, our analyses were based solely on the primary assembly rather than on both the primary and alternate assemblies. Consequently, the data comprise a single haploid genome per individual. This conservative approach was intentionally adopted to ensure high assembly quality, as the alternate assemblies were less contiguous than the primary assemblies.

To evaluate contiguity, we compared our wild mouse assemblies to previously published long-read assemblies from laboratory strains (Ferraj et al. 2023). Our assemblies showed substantial improvement with an average N50 ∼ 20 times larger (130 Mb vs 6.4 Mb), and a greatly reduced L50 (9 vs 132), consistent with longer and fewer contigs (Table 1; Supplementary Table 3).

### Wild mice harbor extensive structural genomic variation

We identified structural variants (SVs) in each genome, relative to the GRCm39 reference genome, and we retained variants with a length of at least 100 bp. Relative to the reference genome, insertions and deletions were roughly equally common and were far more numerous than inversions (Table 1). The average number of SVs in mice from Manaus (65,951) was 16% higher than the average number of SVs in mice from NH-VT (56,684), and this difference was significant (Wilcoxon rank-sum test, *p-value* =. 0.0068; Supplementary Table 4). The total number of SVs among all mice from Manaus (138,392) was also higher than the total number of SVs among all mice from NH-VT (130,898). To assess whether these differences were driven by differences in genome assembly quality, we compared the total number of SVs with various genome assembly metrics. SV count was not associated with scaffold N50, the size of the largest scaffold, read depth, or SVIM score (Supplementary Fig. 1). SV count was significantly associated with genome assembly size, but the difference in SV count between Manaus and NH-VT persisted after normalization by assembly length (two-side Wilcoxon rank-sum test, *p-value*= 0.011). Overall, SVs constitute roughly 160 Mb or 6% of the mouse genome (Table 1).

To compare the number of SVs to the number of single nucleotide mutations, we used DeepVariant (Poplin et al. 2018) to identify single nucleotide variants (SNVs) in all genomes. We identified 19,950,303 SNVs in Manaus and 17,494,606 SNVs in NH-VT. The greater number of SNVs in Manaus compared to NH-VT is roughly proportional to the greater number of SVs in Manaus compared to NH-VT and is consistent with a larger effective population size in Manaus than in NH-VT.

SVs accounted for a substantial fraction of the genomic differences observed among mice within both populations. In Manaus, SVs encompassed 169.2 Mb of the reference genome, compared to 19.9 Mb for SNVs. Thus, SVs comprise over eightfold more DNA sequence than SNVs in Manaus. A similar pattern was observed in NH-VT, where SVs encompassed 159.3 Mb and SNVs encompassed 17.5 Mbp, corresponding to ninefold more sequence for SVs than for SNVs. Although SNVs were numerically more abundant, SVs accounted for the majority of nucleotide differences between genomes in both populations, underscoring the contribution of SVs to genomic diversity.

We used all callable sites to estimate genome-wide nucleotide heterozygosity (π), Waterson’s θ (θ_W_), and Tajima’s D. After filtering SNVs by quality and missing data, we analyzed 2,232,507,882 sites in Manaus and 2,207,831,691 sites in NH-VT. Mean per-site nucleotide diversity was higher in Manaus (π = 0.0017) than in NH-VT (π = 0.0014), and this difference was significant (across all genomic windows, paired Wilcoxon single-rank *p-value* < 2.2e^-16^). Waterson’s θ was also higher in Manaus (θ_W_ = 0.0021) than in NH-VT (θ_W_ = 0.0016; *p-value* < 2.2e^-16^). Tajima’s D was slightly negative in both populations (Manaus Tajima’s D = −0.91; NH-VT Tajima’s D = −0.46), indicating a slight excess of rare variants consistent with a recent population expansion (Agwamba et al. 2026).

### Catalog of transposable elements

To create a library of TEs, we first used RepeatModeler2 (Flynn et al. 2020) which recovered an average of 748 TE consensus sequences per genome, approximately 70% of which were classified as “unknown” due to limited homology with annotated TE families (Supplementary Table 5). TE libraries were then curated using MCHelper (Orozco-Arias et al. 2024) under both the automatic and manual modes. Because of the computational demands of large genomes, curation was performed on assemblies with the highest coverage from three Manaus individuals (FMM215, FMM221and FMM239) and three NH-VT individuals (MPR135, MPR137, MPR144). During the automatic curation step, we recovered an average of 317 TE consensus sequences per genome (Supplementary Table 6). All consensus sequences were then merged from the three individuals in each population to generate a single population-specific TE library, including redundant sequences, resulting in 985 sequences for Manaus and 923 sequences for NH-VT. In the manual curation step, which included structural validation and removal of redundant sequences, we retained 481 consensus sequences for the Manaus library and 340 consensus sequences for the NH-VT library (Table 2).

**Table 2.**
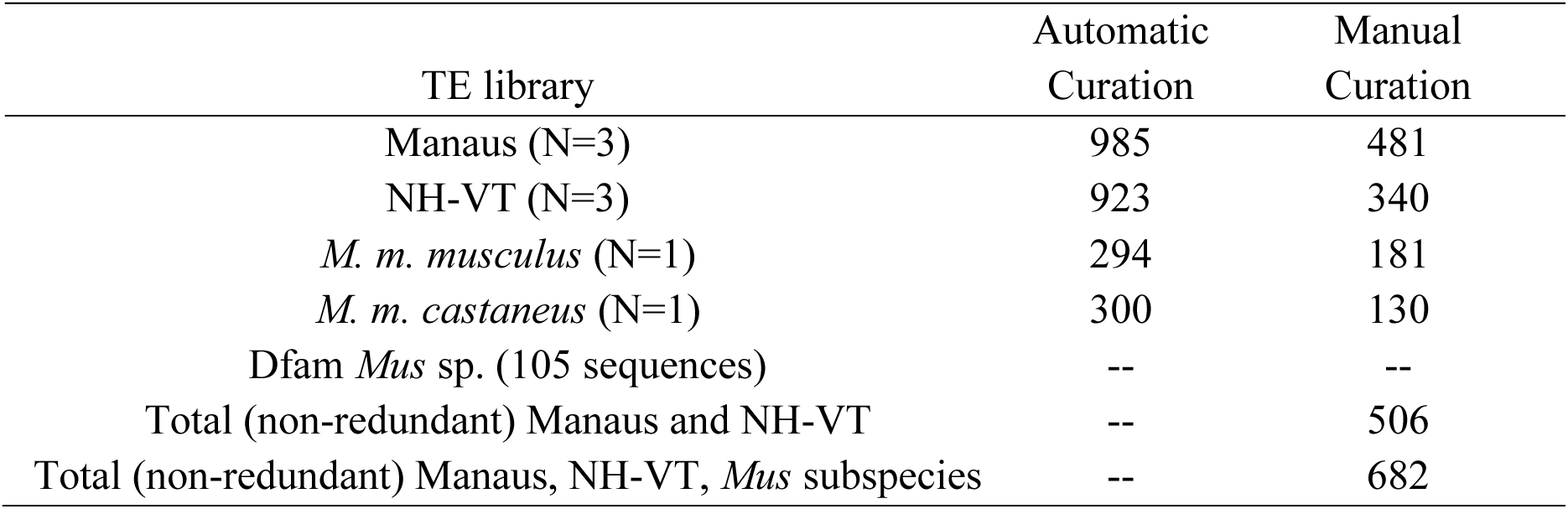
Number of TE consensus sequences following automatic and manual curation steps using MCHelper and Dfam data for *Mus* species.

Finally, all curated TE consensus sequences from both populations were merged, and redundancy was removed using the 80-80-80 rule (Wicker et al. 2007). This resulted in a core set of 506 TE consensus sequences shared between the Manaus and NH-VT populations (Table 2). To further expand the catalog of TEs, we used genome assemblies for *M. m. musculus* (Ferraj et al. 2023) and *M. m. castaneus* (Ferraj et al. 2023), and curated additional TE libraries for both subspecies, recovering 181 and 130 TE consensus sequences, respectively (Table 2). We then combined these libraries with the Manaus + NH-VT TE library, and TE consensus sequences reported for other closely-related *Mus* species in the Dfam database (Hubley et al. 2016; Storer et al. 2021). This final integration resulted a non-redundant TE library containing 682 consensus sequences, including 29 novel TE consensus sequences from LTR (LTR/ERV and LTR/Copia) and LINE/L1 families, with length sizes ranged from 630 bp (LINE/L1) to 10,192 bp (LTR/ERV; Table 2; Electronic Supplementary Material).

### Genome-wide TE annotation

We used the manually-curated TE library to annotate whole-genome assemblies for the 20 wild-caught individuals from Manaus and NH-VT. We also annotated TEs in the genome assemblies of the wild-derived inbred strains SARA and MANA (Supplementary Table 7). Across all annotated genomes, individuals carried on average 1.47 million TE copies, with TEs accounting for approximately 36% of the genome (Supplementary Tables 7-9). The most common TE families were LINES (56.5%), LTRs (30.9%) and SINES (10.3%). Among TE superfamilies, LINE/L1 copies were the most abundant, representing nearly 50% of all TEs, followed by LTR/ERV copies (∼8%) (Fig. 2a).

**Figure 2.**
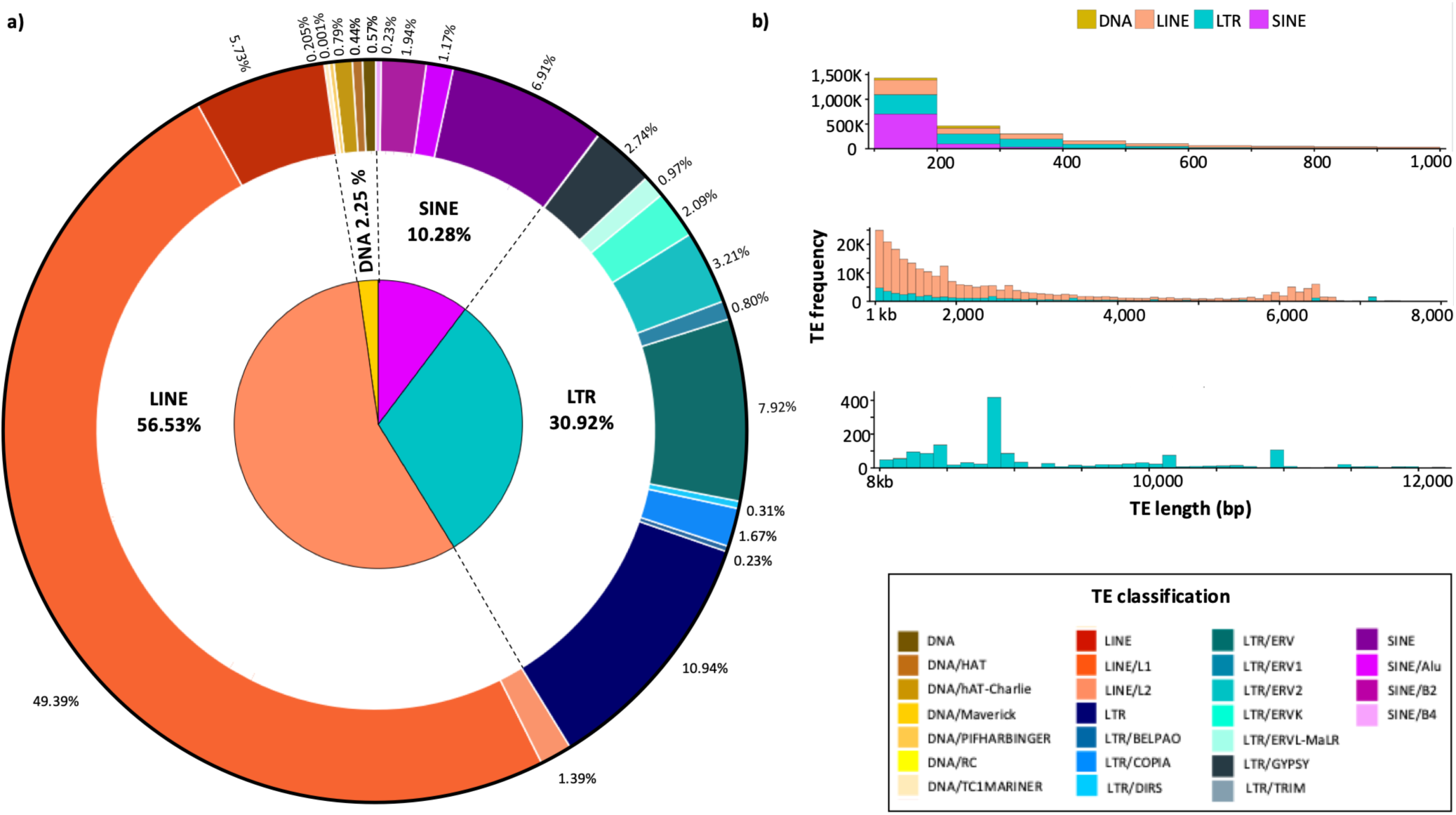
TE landscape in wild-caught mice. a) Average proportion of different TE families and superfamilies annotated across the genomes of 20 wild mice. b) TE length distribution for DNA, LINE, LTR and SINE elements annotated across genomes.

The distributions of TE length were consistent with previous reports in classical inbred mouse strains and other rodent species (Ferraj et al. 2023; Gozashti et al. 2023; Osmanski et al. 2023). For example, SINEs showed characteristic lengths around 200 bp (Fig. 2b), and ERV copies ranged from 480 bp to 11 kb, including some of the longest ERV elements reported in the house mouse genome (Ferraj et al. 2023). TE copies longer than 8 kb were further examined using custom scripts. LTR/ERV and LTR/ERVK were the most abundant TE copies longer than 8 kb, although most represented old, nested insertions (Electronic Supplementary Material). However, some TE copies showing less than 5% divergence from their consensus sequences were identified as complete elements and ranged in length from approximately 8 to 12 kb (Electronic Supplementary Material).

We compared the TEs annotated in this study with those identified in two previous studies on house mice (Ferraj et al. 2023; Nellåker et al. 2012) (Supplementary Fig. 2). Nellåker et al. (2012) used short-read sequences of 18 inbred laboratory strains of *Mus musculus* and the closely related species *Mus spretus*, while Ferraj et al. (2023) used long-read sequences and surveyed 20 lab strains including both classical inbred strains and inbred strains representing the genetically distinct subspecies, *M. m. musculus* and *M. m. castaneus.* Comparisons among these three studies is challenging because of the different sampling strategies and different TE identification and annotation methods used. In particular, we annotated all TEs, including those that were fixed and polymorphic, while Nellåker et al. (2012) and Ferraj et al. (2023) identified polymorphic TEs that were associated with SVs. Thus the total number of TEs reported in this study was an order of magnitude greater than in those studies. When comparing TEs associated with SVs across the three studies, we observed the greatest overlap with Ferraj et al. (2023), with ∼19,000 TEs in common (Supplementary Fig. 2). Nonetheless, Ferraj et al. (2023) identified ∼77,000 TEs not seen in this study, and we identified ∼52,000 TEs not seen in Ferraj et al. (2023). A large fraction of the TEs unique to Ferraj et al. (2023) are attributable to the inclusion of the genetically distinct subspecies *M. m. musculus* and *M. m. castaneus*, while the unique TEs in this study are attributable to the greater diversity in natural populations of wild mice compared to the diversity captured by lab strains. This observation is consistent with previous studies showing that classical inbred strains capture a small amount of the variation segregating in natural populations (e.g. Salcedo et al. 2007; Yang et al. 2011; Dumont et al. 2024). Approximately half of insertion SVs in our study overlapped with TEs (Supplementary Table 4), a result that is broadly consistent with Ferraj et al. (2023) who reported that ∼39% of SVs are attributable to transposable element variants.

### TE variation across environments

To identify polymorphic TE insertions in wild-caught individuals, we mapped flanking regions of each TE in each wild-caught individual to the house mouse reference genome (GRCm39), and then we reciprocally mapped flanking regions of each TE in the reference genome to the genomes of each wild-caught individual, as in Vargas-Chavez et al. (2022) (see Methods for details; Supplementary Fig. 3). We identified roughly 1.29 million euchromatic TE copies in each population (Supplementary Table 10). We then retained those for which reciprocal mapping enabled us to identify the presence or absence of a TE insertion in at least 18 individuals, resulting in a total dataset of 1,127,190 TE copies. The complete dataset with the distribution of all TEs in all 20 individuals is available for download from the Electronic Supplementary Material. Rarefaction analysis showed a rapid saturation of TE family discovery, with accumulation curves flattening after the inclusion of approximately seven genomes, indicating that most TE families are shared among individuals and are captured in our TE annotation and assemblies (Supplementary Fig. 4).

Principal component analysis (PCA) of TE polymorphisms revealed clear population structure, with Manaus and NH-VT individuals clustering separately (Fig. 3a). We estimated TE copy frequency among the total dataset of 1,127,190 annotated euchromatic TEs, and classified insertions as fixed (present in all individuals), common (present in ≥ 10% and ≤ 95% of the individuals), or rare (present in <10% of the individuals). As expected, most insertions were fixed, indicating that most TE copies represent ancient insertions. Across all TEs, 96% of the copies were fixed (1,082,192 TEs), 3.7% of TE copies were common (42,071 TEs), and 0.3% of TE copies were rare (2,927) (Fig. 3b; Supplementary Fig. 5; Supplementary Table 11). Within each category, SINE, LINE/L1, LTR, and LTR/ERV families were the most abundant (Fig. 3c; Supplementary Table 11). Interestingly, LINE/L1, LTR and LTR/ERV families constituted a larger proportion of rare TEs; these families have previously been reported to be active in *M. musculus* sp. (Ferraj et al. 2023). This pattern is consistent with evidence from mice that LINE/L1 and several ERV-related LTR families remain retro-transpositionally active and can generate heritable germline insertion polymorphisms (Gagnier et al. 2019).

**Figure 3.**
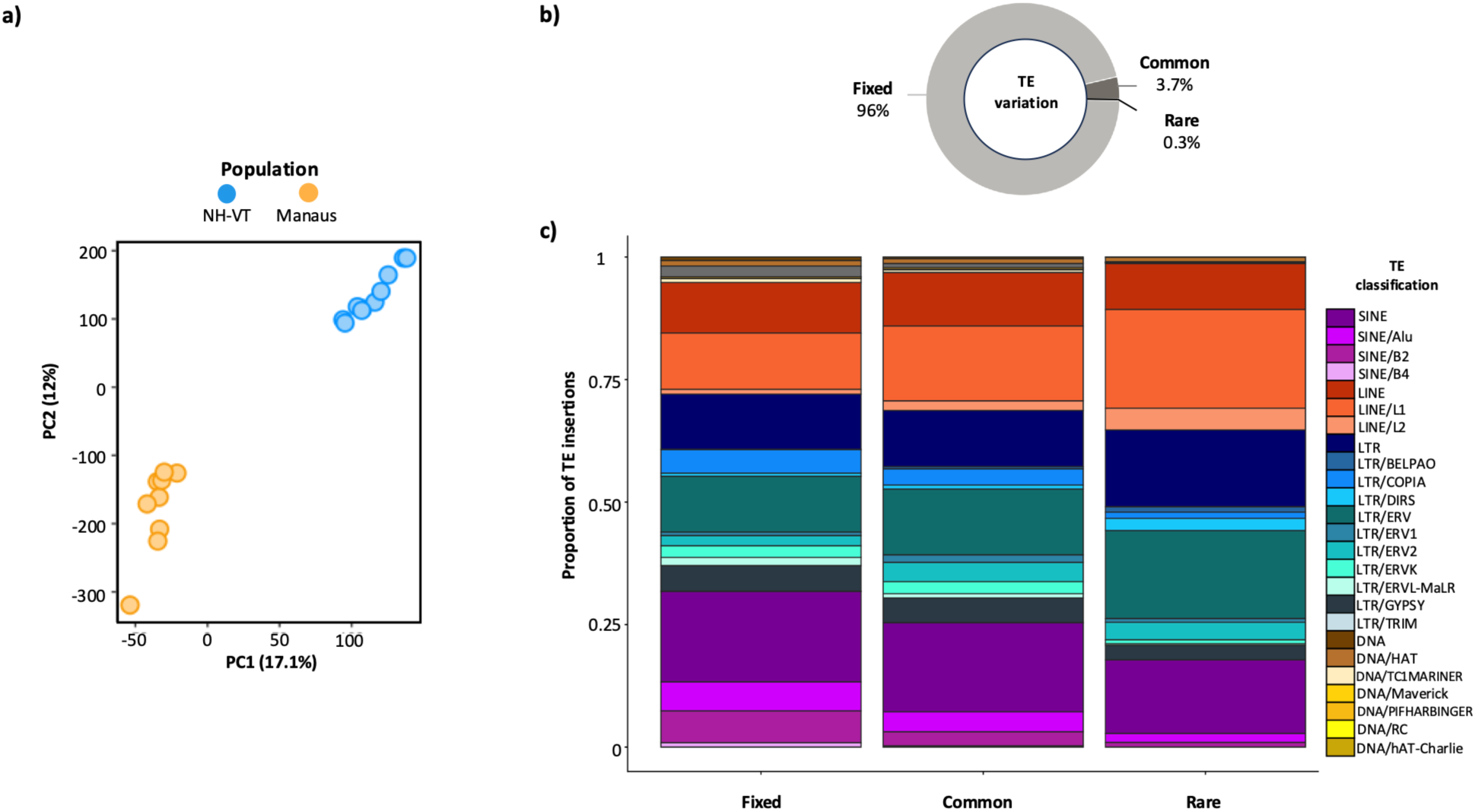
TE insertions in euchromatin regions. a) Principal Component Analysis of all TE insertions in euchromatic regions. b) Proportion of TE insertions that are fixed, common, or rare. c) Proportion of TE family composition across frequency categories.

We next identified high-frequency population-specific insertions (hfTEs) present in ≥ 8 individuals from one population and absent in ≥ 8 individuals from the other population. High-frequency TE insertions were more than twice as common in NH-VT as in Manaus (Supplementary Table 12). In NH-VT, we identified 1,354 hfTEs, dominated by LTRs (41.2%) and SINEs (22.2%; Fig. 4a; Supplementary Tables 12 and 13). The lengths of these TE insertions ranged from 100 bp (DNA and SINE families) to > 6 kb, with the largest insertion being a 9.1 kb LINE element (Supplementary Fig. 6). In Manaus, we identified 573 hfTEs (Supplementary Tables 12 and 14), again dominated by LTR families (18.2%: Fig. 4a). These TE insertions ranged from 100 bp to over 6 kb, with large insertions observed in LINE, LTR, and multiple LTR/ERV families (Supplementary Fig. 6). We observed broadly similar TE family distributions among hfTEs in NH-VT and in Manaus (Figure 4a; Supplementary Table 12).

**Figure 4.**
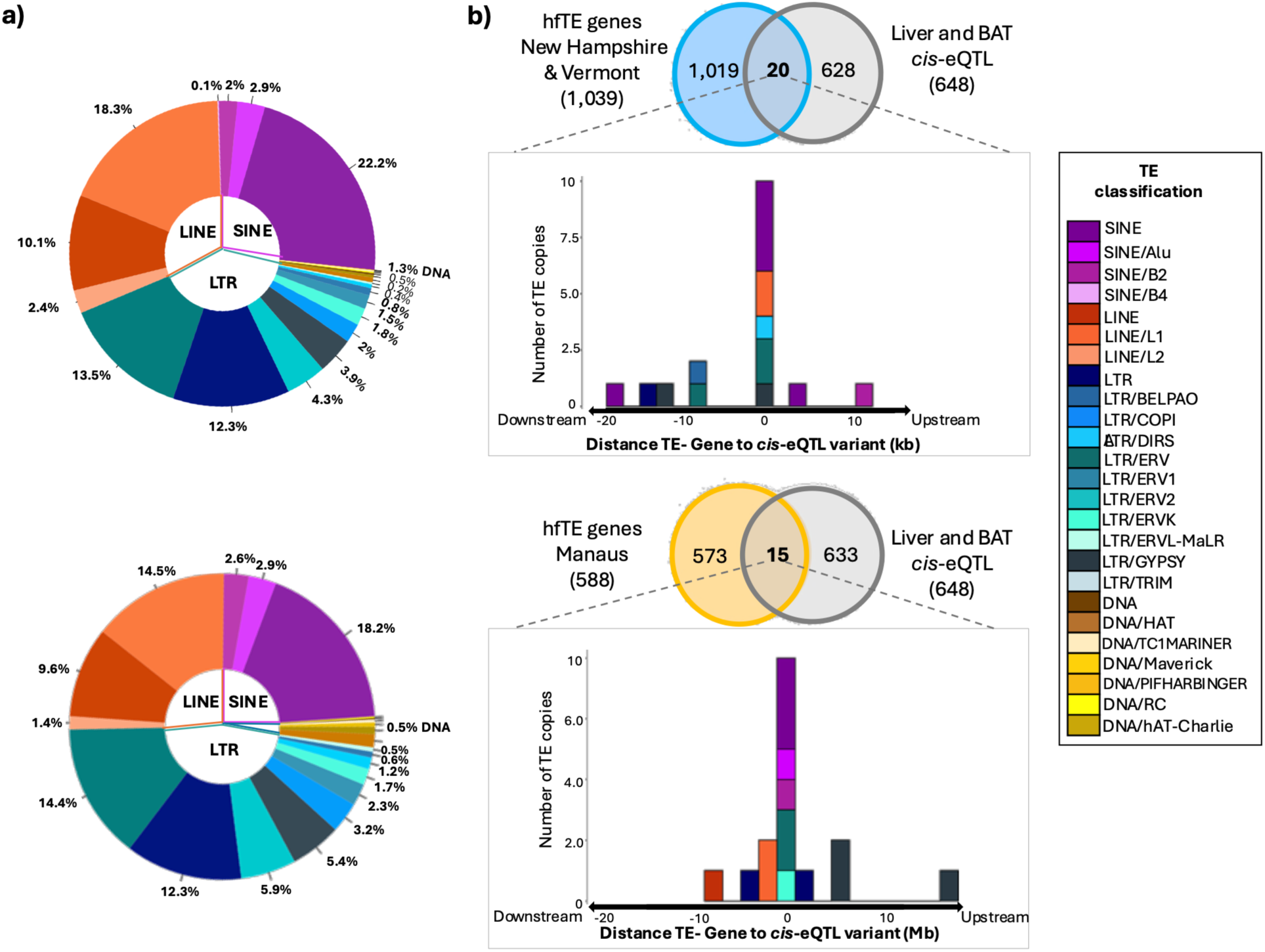
High-frequency population-specific TE insertions (hfTE). a) Family composition of hfTEs in NH-VT (top) and Manaus (bottom). In NH-VT, we identified 1,354 hfTEs; in Manaus, we identified 573 hfTEs. b) Venn diagrams illustrating the overlap between hfTE genes and genes associated with cis-eQTL variants identified in liver and brown adipose tissue from crosses between wild-derived inbred strains of mice (Durkin et al. 2024). In NH-VT, we identified 1,039 overlapping genes, and in Manaus, we identified 588 overlapping genes (see text for details). For overlapping genes, the distance between the TE-gene and the corresponding cis-eQTL variant is shown.

We experimentally validated a subset of hfTEs using PCR amplification of fragments using primers designed to match flanking sequences (see Methods; Supplementary Table 15). We tested five hfTEs in Manaus and seven hfTEs in NH-VT for all 20 individuals. Across 240 amplifications (20 individuals x 12 loci), 230 tests (95.8%) confirmed the predicted presence/absence of a particular insertion (Supplementary Table 15).

### Gene expression variation and TE insertions in wild mice

TE insertions may affect the expression of genes by adding, altering, or disrupting *cis*-regulatory elements (Feschotte 2008). To assess whether TE insertions affect the expression of nearby genes, we generated and analyzed RNAseq data from liver in the wild-caught mice from Manaus and NH-VT. We studied liver because it is an important tissue for growth and metabolism, traits that differ between temperate and tropical mice (Ballinger et al. 2023). Differential expression analysis identified 3,970 differentially expressed genes (DEGs) between populations (*p-value < 0.05*; Supplementary Table 16).

We next looked for overlap between these 3,970 DEGs and the hfTEs in each population. In NH-VT we found that 123 of these DEGs had a hfTE within 20 kb (Supplementary Fig. 7), while in Manaus, we found that 71 of these DEGs had a hfTE within 20 kb (Supplementary Fig. 8). In both cases, the overlap between hfTEs and DEGs was significantly more than expected by chance, indicating that some hfTEs underlie expression differences between tropical and temperate mice [Pearson’s *X^2^* (1,N=17,024)=4.90 in NH-VT and 8.40 in Manaus*, p-value* < 0.027) Supplementary Table 17].

### TE insertions and cis-regulatory expression differences in wild-derived inbred strains

Studying expression variation in wild mice is useful because it captures variation in natural populations (Mack et al. 2018). However, environmentally-induced variation in expression is not controlled in wild mice which may differ in age, reproductive status, and health. Moreover, expression differences in wild mice could be influenced by hfTEs acting in *cis*- or in *trans*-, yet only *cis*-acting hfTEs are expected to lie near the genes whose expression they influence.

To study expression differences in a controlled environment and to identify *cis*-regulated expression quantitative trait loci (*cis*-eQTL), we used expression data from RNA-seq in liver and brown adipose tissue (BAT) in the wild-derived inbred strains SARA and MANA (Durkin et al. 2024). By using crosses between these strains, we were able to identify *cis*-eQTL by studying allele-specific expression (ASE) patterns in F1 mice. Crosses between SARA and MANA identified 648 *cis*-eQTL from ASE (Durkin et al. 2024). To integrate these expression data with patterns of TE insertion polymorphisms, we first asked whether the hfTEs in NH-VT and Manaus were present in the SARA and MANA strains, respectively. Of the 1,354 hfTEs present in NH-VT, 879 were also found in SARA. Of the 573 hfTEs present in Manaus, 502 were also found in MANA.

We then identified all genes within 20 kb of these hfTEs (hfTE genes) (Supplementary Fig. 9). In NH-VT, 699 of the 879 hfTEs (80%) were within 20 kb of one or more genes, corresponding to 1,039 unique genes (Fig. 4b). In Manaus, 418 of the 502 hfTEs (83%) were within 20 kb of one or more genes, corresponding to 588 unique genes. In both populations, individual hfTEs were sometimes associated with multiple genes (e.g. a LINE/L1 insertion fell within 20 kb of *Apol7a*, *Apol9a*, *Gm20688*, *Gm3787*, and *Gm49436*; Supplementary Table 13). In contrast, multiple hfTEs sometimes mapped to the same gene. In some instances, these different insertions represented different TE families (e.g. SINE and LTR-DIRS insertions were associated with *Rabif, Gm15454 and Gm25612*; Supplementary Table 14).

To test whether hfTEs were typically associated with *cis*-regulated gene expression differences between SARA and MANA, we examined the overlap between hfTE genes and genes exhibiting *cis*-eQTL (Durkin et al. 2024). In NH-VT, we found that only 20 of 1,039 hfTE genes overlapped with *cis*-eQTL genes (Fig. 4b). Similarly, in Manaus, only 15 of 588 hfTE genes overlapped with *cis*-eQTL genes (Fig. 4b). In neither case was the overlap more than expected by chance [Pearson’s *X^2^* (1,N=4,555)=0.063 in NH-VT and 3.2 in Manaus*, p-value* > 0.07), Supplementary Table 18]. Nonetheless, a few genes adjacent to hfTEs did show altered *cis*-regulatory expression patterns, and the fact that they are associated with population-specific high-frequency insertions makes them reasonable candidates as targets for selection underlying adaptive differences. Strikingly, nearly all of these genes (33 out of the 35 hfTE genes exhibiting a *cis-eQTL*) were identified in previous studies as candidates for selection based on population branch statistic (PBS) tests (Durkin et al. 2024) or environmental association analysis using latent factor mixed models (LFMM) in wild-caught populations (Phifer-Rixey et al. 2018; Ferris et al. 2021; Gutiérrez-Guerrero et al. 2024).

In both NH-VT and Manaus, most of the hfTE genes exhibiting a *cis*-eQTL corresponded to TEs falling within gene coordinates (primarily intronic regions or genic regions), while others were located in upstream or downstream regions (Fig. 4a-b). Among these, SINE and LTR elements were the most frequent. In NH-VT, the phenotypes associated with the 20 hfTE genes with a *cis*-eQTL were principally related to cholesterol and lipid metabolism, body compositions, glucose tolerance, grip strength, and behavior (Supplementary Table 19). In Manaus, the phenotypes associated with the 15 hfTE genes with a *cis*-eQTL were principally related to body composition, glucose tolerance, and thermoregulation (Supplementary Table 19).

Finally, we looked for overlap between hfTE genes, wild-caught DEGs, and genes harboring a *cis*-eQTL in the SARA-MANA cross. This comparison identified five genes in NH-VT (*Abcc6, Acsm5, Sec14l2, Ano10,* and *Lox*) and four genes in Manaus (*Sypl1*, *Fstl1, Cd3e,* and *Tm7sf3*) (Table 3 and Fig. 5; Supplementary Table 20). The annotated functions of these genes are associated with cholesterol-lipid metabolism and body composition phenotypes (Fig. 5).

**Table 3.**
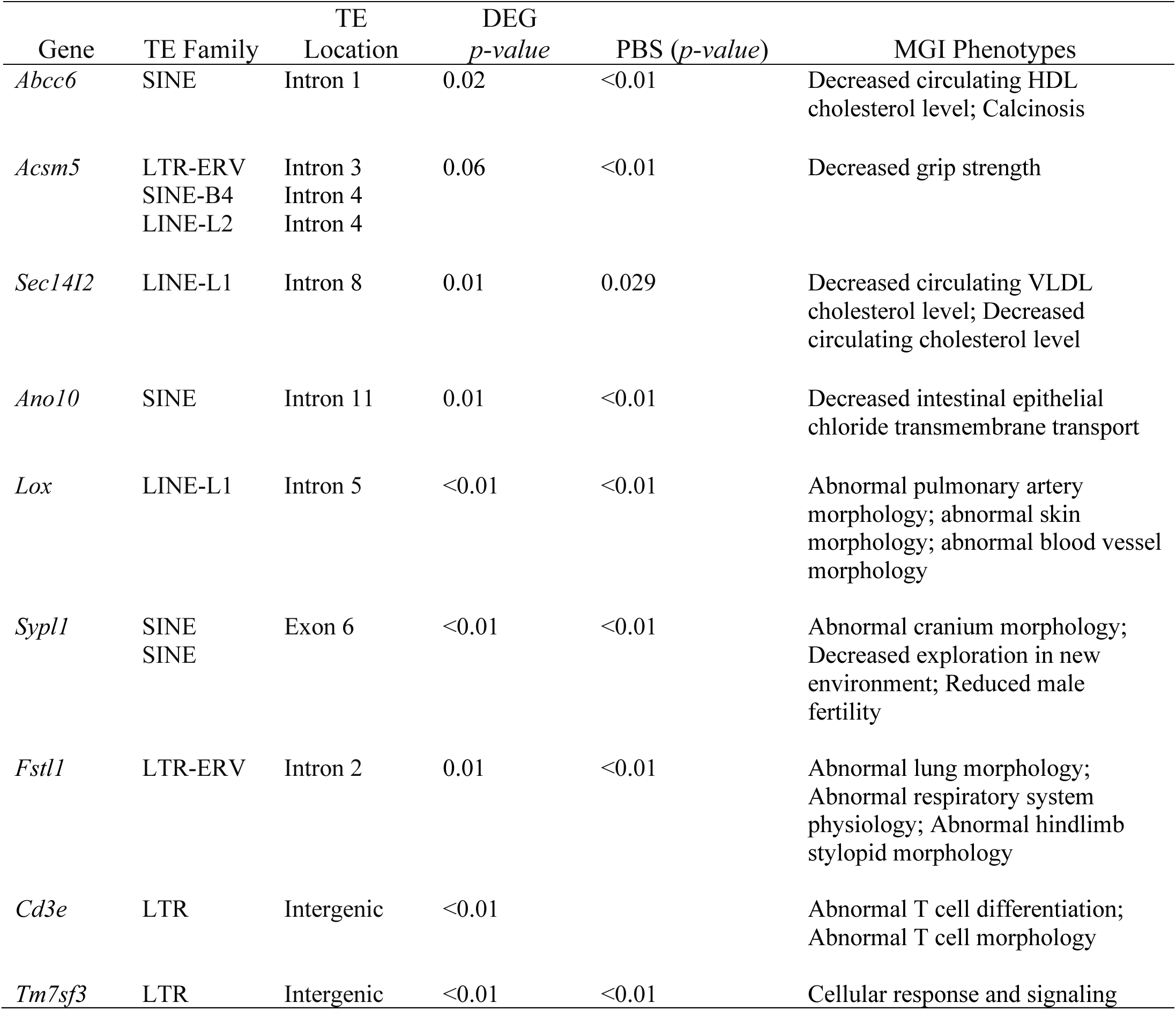
Characterization of hfTE genes associated with a *cis*-eQTL and significant DEG.

**Figure 5.**
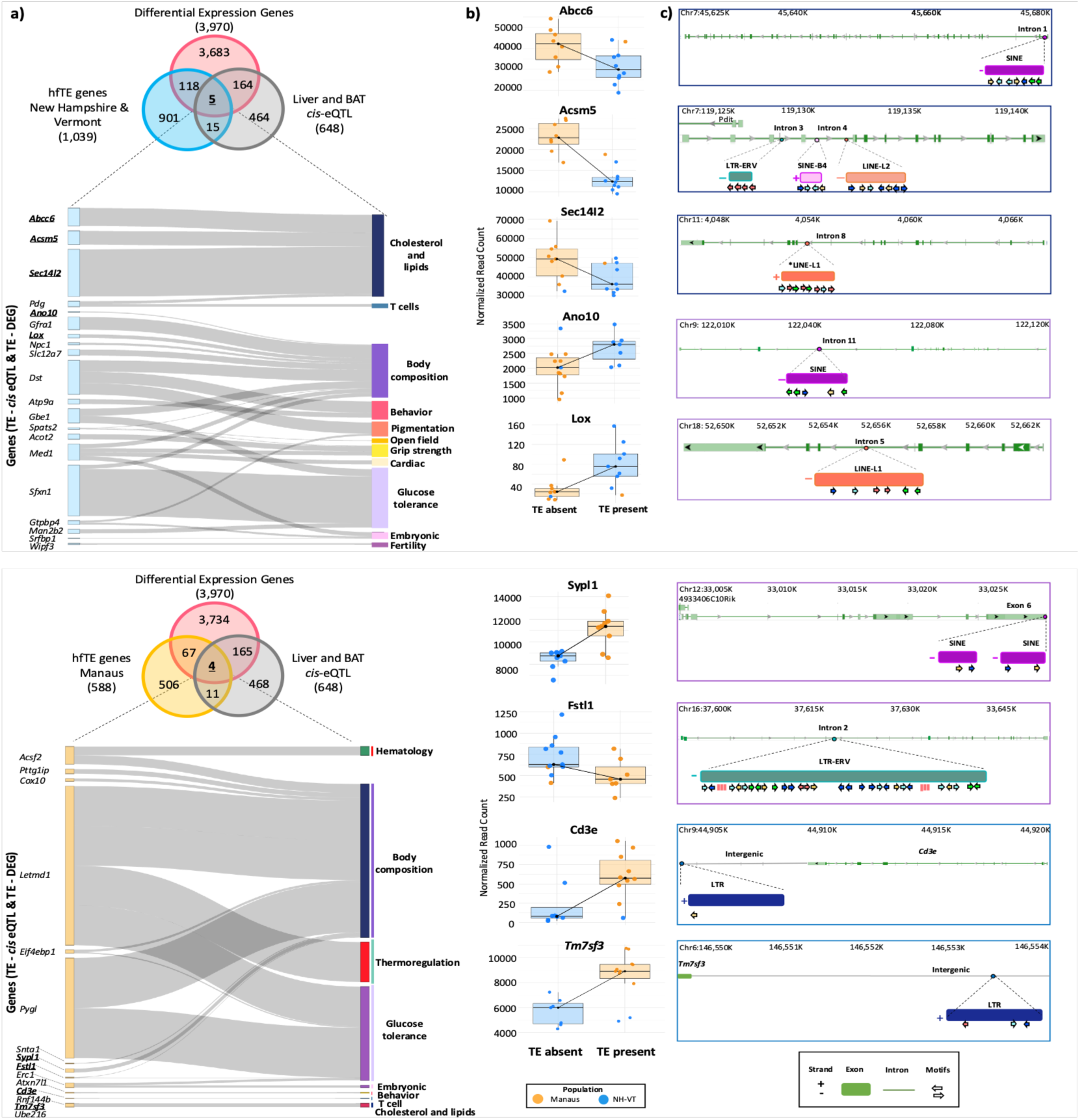
High-frequency TE insertion (hfTE) genes, differentially expressed genes (DEGs) between temperate and tropical wild-caught mice, and *cis*-eQTL identified from crosses between temperate and tropical wild-derived inbred strains. a) Venn diagrams showing the overlap between DEGs, hfTE genes, and genes associated with *cis*-eQTL for NH-VT (top) and Manaus (bottom). The Sankey diagram illustrates the phenotypes reported in the MGI database for the overlapping hfTE genes and *cis*-eQTL genes (20 genes in NH-VT and 15 genes in Manaus). The width of each connection reflects the expression levels reported in Durkin et al. (2024). b) Boxplots showing normalized RNA-seq read counts for genes shared among DEGs, hfTE genes, and *cis*-eQTL genes, grouped by the TE-genotype (present/absent) across individuals. c) Maps of these eight overlapping genes (five in NH-VT, top, and three in Manaus, bottom) showing locations of hfTEs for each gene, together with predicted transcription factor binding motifs identified within the TE sequences.

To further evaluate whether polymorphic TE insertions may participate in host regulatory networks, we identified transcription factor (TF) bindings motifs within hfTEs (Fig. 5c). As expected, zinc-finger TFs were the most prevalent across TE sequences (Zhou et al. 2017). Moreover, in both populations, hfTEs were densely populated by motifs from diverse regulatory families (Supplementary Table 21-22). We observed very strong signatures for canonical regulators of promoters and enhancers (e.g., KLF and SP factors in LINE/L1-*Lox* and LTR/ERV-*Fstl1* TE-genes), along with PATz, VEZF RFX and WT factors, which are known to cooperate at GC-rich *cis-regulatory* modules and chromatin-associated regulatory sites, indicating a densely wired regulatory hub. In addition, we detected TF motifs associated with growth-response regulators (e.g., EGR, SOX, GLI, TWIST, TWST), and numerous regulators involved in hypoxia-responsive muscle, circulatory, vascular, developmental and immune pathways (Supplementary Tables 21-22). These results suggest that polymorphic TE insertions provide regulatory substrates capable of shaping population-specific gene expression responses to environmental pressures.

## Discussion

Recent studies in well-characterized model organisms have documented the landscape of TE insertion polymorphisms and shown that TE variation can influence gene regulation and underlie adaptation. Examples include LTR insertions associated with regulation of the *Flowering locus C* under heat stress in *Arabidopsis* sp. (Raingeval et al. 2024) and TE polymorphisms affecting gene regulation and adaptation to stress conditions in *D. melanogaster* (Ullastres et al. 2021; Merenciano and González 2024). By contrast, although the house mouse is a major vertebrate model system, the landscape of TE variation in natural populations and the contribution of TE variation to adaptation have been comparatively underexplored, largely due to the complex architecture and large genome sizes characteristic of mammals and the computational effort required to resolve repetitive sequences (Osmanski et al. 2023). In this study, long-read sequencing of wild-caught mice from populations in different environments provided new insight into the genetic and regulatory impact of polymorphic TE insertions.

Although long-read sequencing greatly improves resolution of repetitive regions, accurate characterization of TE landscapes still depends on robust computational resources for TE-library curation, annotation and TE polymorphism discovery. Previous studies of TE landscapes in *M. m. domesticus* and close relatives have applied different experimental and computational strategies and different taxa, complicating direct comparison of TE copy number across studies (Nellåker et al. 2012; Ferraj et al. 2023). The greater number of TE variants identified by Ferraj et al. (2023) and Nellåker et al. (2012) among laboratory strains is attributable to the inclusion of different subspecies and species of mice. In contrast, our results reflect variation among individuals within natural populations. Moreover, by using a manually curated TE library, we were able to identify 29 novel TE consensus sequences. Continual refinement of TE annotation frameworks in complex genomes will be essential for understanding the contribution of TEs to genome function and adaptation across environments and physiological contexts.

We found that SVs contribute substantially to genomic diversity in natural populations of mice. Consistent with patterns in humans (Sudmant et al. 2015, Karageorgiou et al. 2024), SVs are numerically less common than SNVs, but they account for a far greater total proportion of the variable bases within natural populations. As such, they are potent source of raw genetic variation on which selection could potentially act. Interestingly, approximately half of all insertion SVs overlapped TEs, indicating that TE-associated insertions represent a major component of structural variation. The presence of low-frequency population-specific insertions suggests that ongoing retrotransposition continues to generate structural diversity, consistent with the known high rate of retrotransposition of LINE/L1 elements in laboratory mice (Richardson et al. 2017). These processes therefore continue to shape genome architecture in natural mouse populations and may contribute to functional divergence.

Neutral or deleterious mutations are not expected to reach high-frequencies in populations. Thus, the hfTEs are reasonable candidates for insertions that may be advantageous. Notably, we observed more than twice as many hfTEs in NH-VT (1,354) as in Manaus (573). This is consistent with previous work showing greater adaptation of house mice to cold environments than to warm environments (Ballinger et al 2023; Durkin et al. 2024). House mice were introduced into the Americas from southern Spain within the last few hundred years. The cold environment of New Hampshire and Vermont is farther from the ancestral climate of house mice in the Iberian peninsula than is the warm climate of Manaus.

We found significant overlap between hfTEs and DEGs, indicating that TE insertions do sometimes affect the expression of nearby genes, consistent with other studies of natural populations (e.g. Rech et al. 2022). By comparing these DEGs with *cis*-eQTL identified in laboratory crosses, we identified a small set of genes that likely contribute to adaptive gene expression differences. These observations are consistent with many studies showing that a large fraction of *cis-*regulatory elements may ultimately derive from TE sequences (Lowe and Haussler, 2012; Sundaram and Wysocka, 2020; Fueyo et al. 2022). Thus, although most insertions are likely to be detrimental, TEs nevertheless represent a meaningful source of regulatory material. Consistent with this view, the small overlapping subset of hfTEs associated with altered expression in the cold-adapted NH-VT population involved genes associated with body composition, glucose tolerance and thermoregulation (*Abcc6, Acsm5, Sec14l2, Ano10* and *Lox*), phenotypes that are likely to be important in adaptation to cold. Notably, all of these genes also showed signatures of positive selection (Durkin et al. 2024; Gutiérrez-Guerrero et al 2024). Some of these TE insertions showed a significant enrichment for promoter and enhancers motifs, including KLF and SP factors, as well as PATz, VEZF RFX and WT binding sites, consistent with densely wired regulatory hubs. Overall, our findings suggest that only a minority of polymorphic TE insertions acquire regulatory functions, but those that do may contribute to functional genetic variation and environmental adaptation in population specific contexts. In this sense, our findings echo the regulatory model of Britten and Davidson (1969), who proposed that TE-derived regulatory modules could participate in gene-regulation networks, providing substrates for regulatory innovation and adaptation.

## Methods

### Samples

Western house mice (*Mus musculus domesticus*) were wild-caught from one population in Manaus, Brazil (N=10) and one population at the border of New Hampshire and Vermont (NH-VT), USA (N=10). The Manaus population is from 3°S latitude, with a mean annual temperature of 27°C and an average low temperature of 23°C (Fick and Hijmans 2017). The NH-VT population is from 44°N latitude, with a mean annual temperature of 9°C and an average low temperature of −12°C. Mice from these different locations have adapted to their respective environments through changes in morphology, physiology, and behavior as described previously (Ballinger et al. 2023, Dumont et al. 2024). Mice were collected with Sherman live traps and sacrificed in accordance with a protocol approved by the Institutional Animal Care and Use Committee (ACUC) of the University of California, Berkeley. Each mouse was collected at least 500 m from every other mouse to avoid sampling relatives. Mice were sacrificed in the field and tissues were flash-frozen in liquid nitrogen or stored on dry-ice and then transferred to a −80°C freezer. All mice have been prepared as museum specimens and deposited in the collections of the UC Berkeley Museum of Vertebrate Zoology as described previously (Phifer-Rixey et al. 2018; Gutierrez et al. 2024) (exact collecting localities, dates, and catalog numbers are given in Supplementary Table 1).

We used published gene expression data from two wild-derived inbred strains of *M. m. domesticus* from these same geographic regions (Durkin et al. 2024). SARA is an inbred strain derived through full-sib mating of wild-caught mice from Saratoga Springs, NY. This location is geographically close to the NH-VT population, and SARA mice are genetically similar to mice from NH-VT (Durkin et al. 2024). MANA is an inbred strain derived through full-sib mating of wild-caught mice from Manaus, Brazil. Both strains have been inbred for more than 20 generations.

### High Molecular Weight DNA extraction

Frozen lung and heart tissues were used for high molecular weight (HMW) gDNA extractions, following the Nanobind HMW DNA extraction protocol (Circulomics DNA kit extraction). Approximately 25 mg of tissue was finely minced into pieces ≤1 mm³ using a scalpel. The minced tissue was transferred to chilled Dounce homogenizers and kept on ice throughout the entire lysis process. The lysate was incubated with Proteinase K at 55°C and then centrifuged at 700 rpm for 3-5 minutes. RNase A buffer was then added, and incubation continued for an additional 30-40 minutes at 55°C. Samples were transferred to new tubes, followed by the addition of one Nanobind disk and isopropanol to the lysate. The samples were mixed on a rotator at 20 rpm for 1 hour. The HMW gDNA pellet was washed three times. After washing, the buffers and ethanol were dried, and the Nanodisk was rehydrated using the PacBio Elution Buffer. This solution was incubated at room temperature overnight to allow the DNA to solubilize. The purity and quantity of HMW gDNA extractions were evaluated with Nanodrop and Qubit. Additionally, gDNA samples showing a broad distribution of DNA (25-100 kb) on Femto Pulse were selected.

### Long-read whole genome sequencing

Libraries were prepared for each individual. The sheared gDNA sample was concentrated with AMPure PB Beads (PacBio), followed by end repair and A-tailing to remove single strands and repair DNA damage. The library was subjected to size selection using the Blue Pippin system (>10 kb). The library was then purified and sequenced on a PacBio REVIO platform. Each sample was sequenced to a minimum of 20-fold coverage, with a HIFI read quality > Q34.

### Long read whole-genome assembly

For each individual sequenced, raw HiFi bam files were converted to FASTQ format. For the SARA wild-derived strain, genomic data in FASTQ format was downloaded from the SRA-NCBI database (project accession number: PRJNA1037121). Adapters and low-quality reads were removed using the script *pbadapterfilt.sh* v.2.0 (Sim et al. 2022). Whole genome assemblies for each individual were constructed with hifiasm (Cheng et al. 2021) and polished with Minimap2 v. 2.26 (Li, 2018) and Racon v. 2 (Vasser et al. 2017) running two polishing iterations. Genome assembly metrics were evaluated and summarized using QUAST (Gurevich et al. 2013) and BUSCO (Simão et al. 2015) with the Glires ortholog database. Additionally, scaffolding was performed using the house mouse reference genome (GRCm39 from NCBI) with RagTag v. 2.1.0. (Alonge et al. 2022). Assembly metrics were evaluated a second time (Table 1). The mitogenome was obtained with MitoHiFi v. 3.01 (Uliano-Silva et al. 2023). Contigs identified as mitogenome were evaluated using a genome alignment with MAFFT (Katoh et al. 2002) using the *M. musculus domesticus* reference mitochondrial genome (NC_005089 from NCBI).

### Structural variant characterization

Structural variants (SVs) were identified relative to the GRCm39 reference genome using SVIM v.2 and Sniffles2 (Heller and Vingron 2020; Smolka et al. 2024). For each individual, bam sequencing files were aligned to the GRCm39 reference genome using Minimap2. Low-quality calls were discarded using the SVIM score ≥ 15 and Sniffles2 coverage >10. We retained SVs with a length ≥ 100 bp. Finally, we merged and identified SVs detected by both methods across all individuals within each population.

### Genetic diversity calculation

PacBio HiFi reads were aligned to the GRCm39 reference genome using pbmm2 (Li et al. 2018) with default parameters and sorting bam files. Variant calling was performed with DeepVariant (Poplin et al. 2018) using --model_type=PACBIO. The resulting VCF files were filtered with bcftools to remove low quality sites and sites with missing data (Danecek et al. 2011). Nucleotide diversity (π), Waterson’s θ (θ_W_), and Tajima’s D were calculated in sliding windows of 10 kb with a 10 kb step using pixy (Korunes and Samuk, 2021).

### TE identification, classification, and library curation

We used RepeatModeler2 (Flynn et al. 2020) to discover and identify TE sequences across the 20 house mice genome assemblies (Supplementary Table 4). We then used MCHelper (Orozco-Arias et al. 2024) to curate the TE library. First, the automatic step of MCHelper was performed using the TE library obtained with RepeatModeler2, along with the Dfam v3.7 database (Storer et al. 2021) and the BUSCO database for Glires. TE characterization and curation with MCHelper were performed on those genomes with the best assembly metrics, including three individuals from Manaus (FMM215, FMM222 and FMM224) and three individuals from NH-VT (MPR134, MPR144 and MPR145), all of which represent the subspecies *M. m. domesticus*. We also included the genomes of the closely related subspecies *M. m. musculus* and *M. m. castaneus* when characterizing and curating TEs since the different subspecies of *M. musculus* are known to share some genetic variation (Phifer-Rixey et al. 2014) (Supplementary Table 4). Subsequently, for Manaus and NH-VT, we concatenated the MCHelper automatic results from the three individuals for each population and removed redundant sequences by applying the 80-80-80 rule (80% identity, 80% coverage and > 80bp; Wicker et al. 2007) with cd-hit (-c 0.80 -aS 0.80; Fu et al. 2012).

To manually curate the TE library, we first used the three individuals from the Manaus population, as well as the genomes of *M. m. castaneus* and *M. m. musculus*, and performed the curation step of MCHelper using the manual inspection option. To detect and remove false positive sequences, such as simple or tandem repeats, we evaluated the TE classification considering the number of full-length copies and the coding domains. Next, to assign already detected families to our consensus sequences, we used BLASTN software (Camacho et al. 2009), applying the 80-80-80 rule, using the Dfam v3.7 database and TE sequences for rodents. To recover high-confident TE consensus sequences that were not passing the 80-80-80 rule, we relaxed the threshold by applying the 70-70 rule and extracted those sequences where percent identity (pid) ≥ 70 and query coverage (qcov) ≥ 70. Finally, for incomplete sequences, we performed BLASTN, using the 80-80-80 rule, and evaluated the classification based on homology using our TE full-length consensus library and Dfam v3.7 database. For the three individuals from NH-VT, we directly applied the 80-80 rule with BLASTN and used the manually curated TE library generated for Manaus and the Dfam v3.7 database as references. After filtering the TE consensus sequences that passed the 80-80 rule, we performed the manual curation steps following the workflow mentioned above. Finally, redundant sequences were removed using cd-hit.

### TE annotation

We annotated each genome using RepeatMasker v. 4.2.1 (Smit and Hubley, 2015) and the curated TE library generated. Later, with the results from this first annotation, we used the OneCodeToFindThemAll Perl tool (Bailly-Bechet et al. 2014) to extend TEs that might be fragmented and generate the final annotation at the family level for each TE element (Supplementary Table 8).

### Heterochromatin masking

Genome assemblies for each individual were subset by chromosome using a custom Python script. The heterochromatin coordinates reported for the house mouse reference genome (GRCm39) were used to identify and mask heterochromatic regions with RepeatMasker v. 4.2.1. Coordinates were obtained from the MGI database based on the MinSat (Guenatri et a., 2004) and ManSat (Packiaraj and Thakur, 2024) chip-arrays. To validate the heterochromatin coordinates, these regions were aligned for each individual with the reference genome using Mummer (Marcais et al. 2018). Once the coordinates were verified, the heterochromatin regions were masked using bedtools maskfasta, and any TE copies overlapping these regions were excluded using bedtools intersect (option -v). Masked regions were visualized using IGV (Robinson et al. 2011) desktop browser and the chip-arrays coordinates.

### TE mapping and TE transfers

To identify polymorphic TE insertions, we mapped 500 bp flanking regions of each TE in each wild-caught individual to the house mouse reference genome (GRCm39), and then we reciprocally mapped flanking regions of each TE in the reference genome to the genomes of each wild-caught individual, as in Vargas-Chavez et al. (2022). Specifically, using the TE annotation files generated by OneCodeToFindThemAll, we generated a bed file for the TEs annotated for each individual. Following Vargas-Chavez et al. (2022), for each TE copy, 500 bp flanking regions (anchors) were generated upstream (F1) and downstream (F2). These anchors were formatted as GFF entries and mapped from the target individual to the reference genome (GRCm39) using the software *Liftoff* v. 1.6.3 (Shumate and Salzberg, 2021), with parameters – polish, --copies, --excluding_partial, and overlap 1, including a polishing step and a copy number tracking option. Only TE copies with successful unique mappings for both F1 and F2 anchors were retained (Supplementary Fig. 3). Mapped anchors were filtered based on (1) the distance between F1 and F2 and length, with insertions ≤ 10 bp tagged as absent (NA), and (2) a TE copy with similar distance (using threshold within 50 bp) between F1-F2 anchors to the original TE length tagged as present. TE copies not meeting these criteria were removed using an R custom script (Supplementary Fig. 3).

BED files were generated for those TEs copies that passed the filters for each target genome. BED files from the 20 individuals were merged and grouped using bedtools groupby, collapsing TEs from the same family that shared similar sizes and insertion coordinates. Grouping was performed using TE family identity and coordinates. Insertions were retained if (1) TE length differed by no more than 50 bp across individuals, (2) at least 50% of the insertion length overlapped, and (3) TE family identity was the same. After merging and filtering, we generated a single bed file containing the mapping information from all the individuals.

To identify TEs that have been mapped across all individuals, consensus TE coordinates were mapped back to each genome using *Liftoff* (with parameters: –polish, –copies, - excluding_partial, and overlap 1). To evaluate if TE copies were present or absent, three filters were considered for the flanking anchors: *location*, *distance* and *length*. TE insertions were considered absent if F1 and F2 anchors were separated by ≤10 bp, and present if the distance approximated the original TE length within ± 100 bp. Both anchors were required to be mapped on the same chromosome or contig (Supplementary Fig. 3). For each TE copy, a binary presence/absence matrix was constructed with rows as TE insertions and columns as individuals. Insertions were coded as 1: present, 0: absent, or NA (missing). Finally, using the coordinates of these TEs, we performed a reciprocal mapping (mapped TE coordinates from the reference genome to individual genomes) to retrieve the TE coordinate for each individual using *Liftoff* (Supplementary Fig. 3).

### TE population-specific polymorphism detection

Using the binary presence/absence matrix previously generated, the frequency of TE copies among individuals was classified into three categories: (1) fixed: TEs present in all individuals; (2) Rare: TEs present in < 10% of the individuals; and (3) Common: TEs present in ≥ 10 % and ≤ 95% of the individuals. To identify population specific TE copies, the frequency of the TE insertions between the Manaus and NH-VT populations were compared. Insertions present in ≥ 80% of one population and absent in ≥ 80% of the other were considered population-specific high-frequency TEs (hfTEs).

The coordinates of hfTEs for each population were used to identify nearby or overlapping genes, using a 20 kb window to account for linkage disequilibrium patterns in house mice (Laurie et al. 2007). Genes located within 20 kb of a hfTE were extracted using bedtools window. A functional enrichment was performed using ShinyGO (Ge et al. 2020), defining as a background the full set of *Mus musculus* genes annotated in the GRCm39 reference genome, and applying a statistical correction (false discovery rate, *p-value* ≤ 0.05).

### Population-specific high-frequency TEs and cis-eQTL variants

We integrated hfTEs with expression data for the wild-derived inbred strains SARA and MANA. Specifically, we used a previously published dataset of 648 *cis-QTL* variants identified in liver and brown adipose tissue from allele-specific expression patterns in F_1_ offspring of SARA x MANA crosses (Durkin et al. 2024). First, we identified hfTEs that were present in the wild-caught individuals as well as in the wild-derived strains (i.e. we identified hfTEs that were present in NH-VT and in SARA, and hfTEs that were present in Manaus and in MANA). For each population, we then identified the set of genes located within 20 kb of a hfTE (hfTE genes). We then identified the coordinates associated with *cis*-QTL variants, using bedtools intersect, and we identified hfTE genes within 20 kb. For each overlapping gene, we also recorded the relative position of the TE insertions (intronic/genic, upstream or downstream) with respect to the gene coordinates.

### RNA extraction and sequencing

The inbred strains SARA and MANA are useful for studying gene expression in a controlled lab environment and for mapping *cis*-eQTL, but they represent only two genotypes and thus may not capture the range of gene expression differences between the populations studied here. To study gene expression more generally in wild mice, we extracted RNA from liver tissues of all 20 wild-caught individuals using the Monarch Spin RNA isolation kit (#T2110, New England Biolabs). Ribodepletion protocol was performed to deplete rRNA from total RNA extracted and to retain only mRNA. Ribodepletion and Library preparation were performed following protocol A from the Watchmaker RNA library Prep Kit (#7BK0002-024, Watchmaker Genomics). The libraries were sequenced using 1 lane of Nova Seq X10B (150bp paired end), generating >63 million raw reads/sample.

### Analysis of RNA-seq data

The raw reads were checked for read quality using FastQC (Andrews, 2010) and quality trimmed with Trimmomatic (Bolger et al. 2014) (parameters: LEADING:3 TRAILING:3 SLIDINGWINDOW:4:15 MINLEN:36). The quality-verified mRNA reads were mapped against the GRCm39 reference genome using STAR with default parameters (Dobin et al. 2013). Two individuals (FMM221 and MPR134) showed extremely low percentages of mapped reads and were therefore excluded from downstream analyses. Gene expression of all the genes was then quantified using FeatureCounts (Liao et al. 2014) and differential gene expression analysis was performed using DESeq2 (Love et al. 2014) in R v. 4.2.2. In DESeq2 we used the design formula: ∼sex + population to identify genes showing differential expression based on population. Benjamini-Hochberg false discovery rate-adjusted P_adj_ < 0.05 was used as a criterion for identifying differentially expressed genes between samples from NH-VT and Manaus.

### Predicted transcription factor binding site discovery

For each hfTE, the corresponding sequence was extracted from individual genome assemblies using samtools faidx. Transcription factor binding sites (TFBS) motifs were identified within TE sequences using the FIMO command-line tool from the MEME Suite (Bailey et al. 2009; Grant et al. 2011), employing the HOCOMOCO v13 database of GRCm39 transcription factor binding profiles (Vorontsov et al. 2024). Motif searches were performed using default parameters. Predicted TRBS were subsequently filtered based on motif significance, retaining only matches with a false discovery rate-corrected q-value < 0.05, as determined from position weight matrix scores.

### PCR validation of TE insertions

We experimentally validated 12 inferred hfTEs using PCR and agarose gel electrophoresis. We first located each candidate TE in the GRCm39 reference genome using UCSC genome browser and retrieved the surrounding DNA sequence (∼650 base pairs upstream and downstream) to provide sufficient context for primer design. Within this region, we annotated the TE, marked the 150 bp flanking regions as the “PCR Target Region", and defined the 500 bp regions beyond that on either side as the “Primer Design Region”. Using Primer3, primers were designed to amplify the “Target Region”, thus amplifying a minimum of ∼300bp PCR fragment. The primer pairs were then verified for their specificity by checking for potential off-target amplification using an UCSC in-silico PCR tool. PCR reactions (50ul) were performed using the OneTaq Master Mix (#M0486, New England Biolabs) under the following conditions: initial denaturation at 94°C for 3 min; 35 cycles of 94°C for 30s, locus-specific annealing temperature for 30s, 68°C for 60s; followed by a final extension at 68°C for 5 min where locus-specific annealing temperature was calculated using the New England Biolab Tm Calculator (https://tmcalculator.neb.com/#!/main). Approximately 6ul of PCR products were run on a 1.2% agarose gel in TBE buffer, stained with Ethidium Bromide and visualized under UV illumination.

TE presence was inferred from the expected size differences between alleles with and without the insertion: a single smaller band indicated absence, a single larger band indicated homozygous presence, and two bands indicated heterozygosity. We tested 12 hfTEs, including five in Manaus and seven in NH-VT. For the five hfTEs from Manaus, all showed the expected size distribution for all 20 individuals. For the seven hfTEs from NH-VT, six showed the expected size distribution for all 20 individuals and one showed the expected size for only 10 individuals (i.e. for one locus an insertion was observed in all 20 individuals but predicted in only 10). Thus across the 20 individuals and 12 loci tested (240 tests), 230 tests (95.8%) experimentally confirmed the predicted presence/absence of a particular insertion.

## Data Access

PacBio HiFi reads sequencing are available at the Sequence Read Archive (SRA) from NCBI and whole-genome assemblies, BioProject accession PRJNA1109785. Supplementary Table 1 includes the complete metadata of the samples: locality information, sex, read sequencing information, and the Museum of Vertebrate Zoology vouchered specimen number and accession. The code and scripts used for the analysis are available from GitHub (https://github.com/YocelynG/Transposable-elements-and-structural-variation). Structural variants bed, along with transposon elements annotations and TE reciprocal mapping are available in Electronic Supplementary Material (DRYAD: 10.5061/dryad.612jm64m9). *Reviewer sharing links*: https://dataview.ncbi.nlm.nih.gov/object/PRJNA1109785?reviewer=3ftdufea4hcncd3mbtm4arpb2r

http://datadryad.org/share/LINK_NOT_FOR_PUBLICATION/pCw2CNJglk04gaCurLb4WGkIgk8rwljZw3lfDSJ8Neo

## Competing of Interest

The authors have declared that no competing interest exist.

## Acknowledgments

We thank the members of the Nachman Lab for their valuable comments and discussions. We thank Lydia Smith, Noëlle Bitter for their technical support and expertise. We thank the members of the Gonzalez Lab for discussions and code assistance. This work was facilitated by access to the SAVIO, UC-Berkeley, USA, Patung cluster, LANCIS-Instituto de Ecología, UNAM; and IPS-CNR, Italy). This research was supported by NIH grants to M.W.N. (R01 GM074245, R01 GM127468, and R35 GM149304). JG was supported by grant EUR2025-164826 funded by MICIU/AEI /10.13039/501100011033 and by grant PID2023-148838NB-I0 funded by MICIU/AEI /10.13039/501100011033 and by FEDER, EU.

## Author contributions

YTGG, JG, and MWN designed the study. YTGG generated the sequence data for all wild-caught mice, JL generated the genome sequencing for MANA, and AV generated and analyzed the RNA-seq data. YTGG conducted most of the analyses, with contributions from SOA, MCZ, JG, and MWN. YTGG and MWN wrote the first draft of the paper, and all authors contributed to subsequent drafts.

## Supplementary Tables

**Supplementary Table 1**. Metadata for wild-caught house mice from Manaus and New Hampshire and Vermont (NH-VT) populations.

**Supplementary Table 2**. PacBio HiFi REvio long-read sequencing summary statistics.

**Supplementary Table 3**. Genome assembly metrics and BUSCO completeness results.

**Supplementary Table 4**. Structural variants (SVs) identified using SVIM and Sniffles relative to the mouse reference genome (Mm39), filtered by quality and length > 100 bp.

**Supplementary Table 5**. Transposable element identification and annotation using RepeatModeler.

**Supplementary Table 6**. TE library generated using the MCHelper automatic step.

**Supplementary Table 7**. Genome-wide annotation for TEs using RepeatMasker.

**Supplementary Table 8**. Total TE counts by family after TE extension.

**Supplementary Table 9**. Proportions of different kinds of TEs across assembled genomes.

**Supplementary Table 10**. Number of TEs in euchromatin regions for each assembled genome.

**Supplementary Table 11**. TE frequencies.

**Supplementary Table 12**. Population-specific high-frequency TEs (hfTEs) in Manaus and NH-VT. hfTEs are defined as being present in ≥ 8 individuals from one population and absent in ≥ 8 individuals from the other population.

**Supplementary Table 13**. hfTEs (presence/absence) in NH-VT.

**Supplementary Table 14**. hfTEs (presence/absence) in Manaus.

**Supplementary Table 15**. PCR validation of candidate TE polymorphic insertions

**Supplementary Table 16**. Differential expression genes identified between Manaus and NH-VT populations.

**Supplementary Table 17**. Overlap between hfTE-genes and differentially expressed genes (DEGs), including permutation test results in Manaus and NH-VT populations.

**Supplementary Table 18**. Overlap between hfTE-genes and cis-eQTL genes, including permutation tests in Manaus and NH-VT populations.

**Supplementary Table 19**. MGI enrichment analysis of genes overlapping hfTEs and cis-eQTL genes (identified in Durkin et al. 2024).

**Supplementary Table 20**. Three way overlap among hfTE-genes, cis-eQTL and DEGs.

**Supplementary Table 21**. Transcription factor motifs in hfTEs in the NH-VT population.

**Supplementary Table 22**. Transcription factor motifs in hfTEs in the Manaus population.

## Supplementary Figures

**Supplementary Fig. 1**. Relationship between structural variant counts and sequencing and assembly metrics. Each point represents one genome with individuals grouped by population: Manaus and New Hampshire and Vermont (NH-VT)). Each panel reports the overall Pearson correlation coefficient and associated *p-value*, together with the within-population correlation.

**Supplementary Fig. 2**. Comparisons of TEs annotated in this study with those in Ferraj et al. (2023) and Nellåker et al. (2012) a). TE counts in different inbred strains as well is in population samples from NH-VT and Manaus. Dotted lines indicate data from Ferraj et al. (2023). b) Top: Venn diagram showing overlap of all annotated TEs. Bottom: Venn diagram showing overlap of polymorphic TEs associated with SVs.

**Supplementary Fig. 3**. Workflow implemented for TE polymorphism identification.

**Supplementary Fig. 4**. Rarefaction analysis showing cumulative curves of annotated TE families across genomes. Rarefaction curves were calculated using python custom scripts available at GitHub (https://github.com/YocelynG/Transposable-elements-and-structural-variation).

**Supplementary Fig. 5**. Frequency distribution of TEs among all 20 mice (top), among 10 mice from Manaus (middle), and among 10 mice from NH-VT (bottom).

**Supplementary Fig. 6**. Length distributions of hfTEs in Manaus and NH-VT.

**Supplementary Fig. 7**. Normalized read counts of the 142 DEGs that overlap with the hfTEs present in NH-VT. Each boxplot shows one hfTE with read counts corresponding to presence (1) or absence (0) of the insertion. Note that some genes are associated with more than one hfTE. Individuals are colored by population of origin: NH-VT (blue) and Manaus (orange).

**Supplementary Fig. 8**. Normalized read counts of the 70 DEGs that overlap with the hfTEs present in Manaus. Each boxplot shows one hfTE with read counts corresponding to presence (1) or absence (0) of the insertion. Note that some genes are associated with more than one hfTE. Individuals are colored by population of origin: NH-VT (blue) and Manaus (orange).

**Supplementary Fig. 9**. Hypothetical physical map of a genomic region showing location of genes, genes with a *cis*-eQTL, TEs, and hfTEs. We identified genes such as A that were associated with a *cis*-eQTL and an hfTE within 20 kb. We excluded genes such as B (with a *cis*-eQTL but no hfTE within 20kb) and C (with an hfTE but no *cis*-eQTL within 20 kb).

**Electronic Supplementary Material (DRYAD,** DOI: 10.5061/dryad.612jm64m9)

-Genome assemblies (scaffolding version)

-Structural Variant Bed files

-TE consensus library (fasta file)

-TE annotations generated with RepeatMasker and OneCode

-Table summary with TE copies with length sizes > 8 kbp

-Reciprocal mapping matrix

-TE insertions overlapping structural variants

## Notes

### Competing Interest Statement

The authors have declared no competing interest.

## References

1. Agwamba KD, Nachman MW. 2023. The demographic history of house mice (*Mus musculus domesticus*) in eastern North America. G3 13(2): jkac332. 10.1093/g3journal/jkac332

2. Agwamba KD, Gabriel SI, Searle JB, Wall J, Nachman MW. 2026. New country for old mice: the recent colonization history of *Mus musculus domesticus* in the Americas. Genome Biol. Evol, in press.

3. Andrews S. 2010. FastQC: A quality control tool for high throughput sequence data. Babraham Bioinformatics. https://www.bioinformatics.babraham.ac.uk/projects/fastqc/

4. Alonge M, Lebeigle L, Kirsche M, Jenike K, Ou S, Aganezov S, Wang X, Lippman ZB, Schatz MC, Soyk S. 2022. Automated assembly scaffolding using RagTag elevates a new tomato system for high-throughput genome editing. Genome Biol 23:258. 10.1186/s13059-022-02823-7

5. Bailly-Bechet M, Haudry A, Lerat E. 2014. One code to find them all: a perl tool to conveniently parse RepeatMasker output files. Mobile DNA 5:13. 10.1186/1759-8753-5-13

6. Bailey TL, Boden M, Buske FA, Frith M, Grant CE, Clementi L, Ren J, Li WW, Noble WS. 2009. MEME SUITE: tools for motif discovery and searching. Nucleic Acids Res 37: W202–W208. 10.1093/nar/gkp335

7. Ballinger MA, Nachman MW. 2022. The contribution of genetic and environmental effects to Bergmann’s rule and Allen’s rule in house mice. Am Nat 199: 691–704.

8. Ballinger MA, Mack KL, Durkin SM, Riddell EA, Nachman MW. 2023 Environmentally robust *cis*-regulatory changes underlie rapid climatic adaptation. Proc Nat Acad Sci USA 10.1073/pnas.2214614120

9. Batzer MA, Deininger PL. 2002. Alu repeats and human genomic diversity. Nat Rev Genet 3(5):370–379. 10.1038/nrg798

10. Bolger AM, Lohse M, Usadel B. 2014. Trimmomatic: A flexible trimmer for Illumina sequence data. Bioinformatics 30(15): 2114–2120. 10.1093/bioinformatics/btu170

11. Bourque G, Burns HK, Gehring M. 2018. Ten things you should know about transposable elements. Genome Biol 19(1):199. 10.1186/s13059-018-1577-z

12. Boursot P, Auffray J-C, Britton-Davidian J, Bonhomme F. 1993. The evolution of house mice. Annu Rev Ecol Syst 24:119–152. 10.1146/annurev.es.24.110193.001003

13. Britten RJ, Davidson EH. 1969. Gene regulation for higher cells: a theory. Science 3891:349–57. 10.1126/science.165.3891.349

14. Brookfield JFY. 2004. Evolutionary genetics: Hiding behind selection. Curr Biol 14(15): R600–R602. 10.1016/j.cub.2004.07.027

15. Camacho C, Coulouris G, Avagyan V, Ma N, Papadopoulos J, Bealer K, Madden TL. 2009. BLAST+: architecture and applications. BMC Bioinformatics 10:421. 10.1186/1471-2105-10-421

16. Casacuberta E, González J. 2013 The impact of transposable elements in environmental adaptation. Mol Ecol 22:1503–1517. 10.1111/mec.12170

17. Charlesworth B, Langley CH. 1989. The population genetics of transposable elements in Drosophila. Genet Res 53(2):139–156. 10.1017/S0016672300023429

18. Cheng H, Concepcion GT, Feng X, Zhang H, Li H. 2021. Haplotype-resolved de novo assembly using phased assembly graphs with hifiasm. Nat Methods 18:170–175. 10.1038/s41592-020-01056-5

19. Coronado-Zamora M, González J. 2023. Transposons contribute to the functional diversification of the head, gut, and ovary transcriptomes across *Drosophila* natural strains. Genome Res 33(9):1541–1553. 10.1101/gr.277565.122

20. Daborn PJ, Yen JL, Bogwitz MR, Le Goff G, Feil E, Jeffers S, Tijet N, Perry T, Heckel D, Batterham P, Feyereisen R, Wilson TG, ffrench-Constant RH. 2002. A single P450 allele associated with insecticide resistance in Drosophila. Science 297(5590), 2253–2256. 10.1126/science.1074170

21. Danecek P, Auton A, Abecasis G, Albers CA, Banks E, DePristo MA, Handsaker RE, Lunter G, Marth GT, Sherry ST. et al. 2011. The variant call format and VCFtools. Bioinformatics 27(5), 2156–2158. 10.1093/bioinformatics/btr330

22. Dobin A, Davis CA, Schlesinger F, Drenkow J, Zaleski C, Jha S, Batut P, Chaisson M, Gingeras TR. 2013. STAR: Ultrafast universal RNA-seq aligner. Bioinformatics 29(1):15–21. 10.1093/bioinformatics/bts635

23. Dumont BL, Gatti DM, Ballinger MA, Lin D, Phifer-Rixey M, Sheehan MJ, Suzuki TA, Wooldridge LK, Frempong HO, Lawal RA. et al. 2024. Into the Wild: A novel wild-derived inbred strain resource expands the genomic and phenotypic diversity of laboratory mouse models. PLoS Genet 20(4):e1011228. 10.1371/journal.pgen.1011228

24. Durkin SM, Ballinger MA, Nachman MW. 2024. Tissue-specific and *cis*-regulatory changes underlie parallel, adaptive gene expression evolution in house mice. PLoS Genet 20(2): e1010892. 10.1371/journal.pgen.1011213

25. Ellison CE, Bachtrog D. 2012. Non-allelic gene conversion enables rapid evolutionary change at multiple regulatory sites encoded by transposable elements. eLife 1:e00176. 10.7554/eLife.00176

26. Ferraj A, Audano PA, Balachandran P, Czechanski A, Flores JI, Radecki AA, Mosur V, Gordon DS, Walawalkar IA, Eichler EE, Reinholdt LG, Beck CR. 2023. Resolution of structural variation in diverse mouse genomes reveals chromatin remodeling due to transposable elements. Cell Genom 3(5):100291. 10.1016/j.xgen.2023.100291

27. Ferris KG, Chavez AS, Suzuki TA, Beckman EJ, Phifer-Rixey M, Bi K, Nachman MW. 2021. The genomics of rapid climatic adaptation and parallel evolution in North American house mice. PLoS Genet 17(4): e1009495. 10.1371/journal.pgen.1009495

28. Feschotte C. 2008. Transposable elements and the evolution of regulatory networks. Nat Rev Genet 9:397–405. 10.1038/nrg2337

29. Feschotte C. 2026. Transposable elements as catalysts of evolutionary innovation. Nat Rev Genet 10.1038/s41576-026-00980-0

30. Fick SE, Hijmans RJ. 2017. WorldClim2: new 1km spatial resolution climate surfaces for global land areas. Int J Climatol 37(12):4302–4315

31. Flynn JM, Hubley R, Goubert C, Rosen J, Clark AG, Feschotte C, Smit AF. 2020. RepeatModeler2 for automated genomic discovery of transposable element families. PNAS 117(17):9451–9457. 10.1073/pnas.1921046117

32. Fu L, Niu B, Zhu Z, Wu S, Li W. 2012. CD-HIT: accelerated for clustering the next-generation sequencing data. Bioinformatics 28(23): 3150–3152. 10.1093/bioinformatics/bts565

33. Fueyo R, Judd J, Feschotte C, Wysocka J. 2022. Roles of transposable elements in the regulation of mammalian transcription. Nat Rev Mol Cell Biol 23(7):481–497. 10.1038/s41580-022-00457-y.

34. Gagnier L, Belancio VP, Mager DL. 2019. Mouse germ line mutations due to retrotransposon insertions. Mobile DNA 10(15). 10.1186/s13100-019-0157-4

35. Galbraith DW, Hayward A. 2023. Transposable elements as regulators of gene expression. Trends Genet 39(6):421–435. 10.1016/j.tig.2023.02.004

36. Ge SX, Jung D, Yao R. 2020. ShinyGO: a graphical gene-set enrichment tool for animals and plants. Bioinformatics 36(8):2628–2629. 10.1093/bioinformatics/btz931

37. Guio L, Vieira C, González J. 2018. Stress affects the epigenetic marks added by natural transposable element insertions in *Drosophila melanogaster*. Sci Rep 8:12197. 10.1038/s41598-018-30491-w

38. González J, Macpherson JM, Petrov DA. 2008. A recent adaptive transposable element insertion near highly conserved developmental loci in *Drosophila melanogaster*. Mol Biol Evol 25(9):1879–1891. 10.1093/molbev/msn143

39. Gozashti L, Feschotte C, Hoekstra HE. 2023. Transposable Element Interactions Shape the Ecology of the Deer Mouse Genome. Mol Biol Evol 40(4): msad069. 10.1093/molbev/msad069

40. Grant CE, Bailey TL, Noble WS. 2011. FIMO: scanning for occurrences of a given motif. Bioinformatics 27(7):1017–8. 10.1093/bioinformatics/btr064.

41. Guenatri M, Bailly D, Maison C, Almouzni G. 2004. Mouse centric and pericentric satellite repeats form distinct functional heterochromatin. J Cell Biol 166(4):493–505. 10.1083/jcb.200403109

42. Gurevich A, Saveliev V, Vyahhi N, Tesler G. 2013. QUAST: quality assessment tool for genome assemblies. Bioinformatics 29(8):1072–1075. 10.1093/bioinformatics/btt086

43. Gutiérrez-Guerrero YT, Phifer-Rixey M, and Nachman MW. 2024. Across two continents: the genomic basis of environmental adaptation in house mice (*Mus musculus domesticus*) from the Americas. PLoS Genet 20(7): e1011036. 10.1101/2023.10.30.564674

44. Hayward A, Clément G. 2022. Transposable elements. Curr biol 32: R904–R909. 10.1016/j.cub.2022.07.044

45. Heller D, Vingron M. 2020. SVIM-asm: structural variant detection from haploid and diploid genome assemblies. Bioinformatics 36(22–23): 5519–5521. 10.1093/bioinformatics/btaa1034

46. Hubley R, Finn RD, Clements J, Eddy SR, Jones TA, Bao W, Smit AF, Wheeler TJ. 2016. The Dfam database of repetitive DNA families. Nucleic Acids Res 44(D1):D81–9. 10.1093/nar/gkv1272

47. Karageorgiou C, Gokcumen O, Dennis MY. 2024 Deciphering the role of structural variation in human evolution: a functional perspective. Curr Opin Genet Dev 88:102240. 10.1016/j.gde.2024.102240

48. Katoh K, Misawa K, Kuma K, Miyata T. 2002. MAFFT: a novel method for rapid multiple sequence alignment based on fast Fourier transform. Nucleic Acids Res 30(14):3059–3066. 10.1093/nar/gkf436

49. Korunes KL, Samuk K. 2021. pixy: Unbiased estimation of nucleotide diversity and divergence in the presence of missing data. Mol Ecol Res 21(4):1359–1368. 10.1111/1755-0998.13326

50. Laurie CC, Nickerson DA, Weir BS, Livingston RJ, Dean MD, Smith K, Schadt EE, Nachman MW. 2007. Linkage disequilibrium in a natural population of the house mouse, *Mus musculus domesticus*. PLoS Genet 3:1487–1495.

51. Liao Y, Smyth GK, Shi W. 2014. featureCounts: An efficient general-purpose read summarization program. Bioinformatics 30(7):923–930. 10.1093/bioinformatics/btt656

52. Li H. 2018. Minimap2: pairwise alignment for nucleotide sequences. Bioinformatics 34(18): 3094–3100. 10.1093/bioinformatics/bty191

53. Love MI, Huber W, Anders S. 2014. Moderated estimation of fold change and dispersion for RNA-seq data with DESeq2. Genome Biol 15:550. 10.1186/s13059-014-0550-8

54. Lowe CB. Haussler D. 2012. 29 Mammalian Genomes Reveal Novel Exaptations of Mobile Elements for Likely Regulatory Functions in the Human Genome. PLoS ONE 7(8): e43128. 10.1371/journal.pone.0043128

55. Lynch CB. 1992. Clinal variation in cold adaptation in mus domesticus: verification of predictions from laboratory populations. Am Nat 139(6):1219–36.

56. Marçais G, Delcher AL, Phillippy AM, Coston R, Salzberg SL, Zimin A. 2018. MUMmer4: A fast and versatile genome alignment system. PLoS Comput Biol 14(1): e1005944. 10.1371/journal.pcbi.1005944

57. Merenciano M, González J. 2023. The Interplay Between Developmental Stage and Environment Underlies the Adaptive Effect of a Natural Transposable Element Insertion. Mol Biol Evol 40(3). 10.1093/molbev/msad044

58. Nellåker C, Keane TM, Yalcin B, Wong K, Agam A, Belgard TG, Flint J, Adams DJ, Frankel WN, Ponting CP. 2012. The genomic landscape shaped by selection on transposable elements across 18 mouse strains. Genome Biol 13, R45. 10.1186/gb-2012-13-6-r45

59. Orozco-Arias S, Sierra P, Durbin R, Gonzalez J. 2024. MCHelper automatically curates transposable element libraries across species. Genome Res 34:1–13. 10.1101/gr.278821.123

60. Osmanski AB, Paulat NS, Korstian J, Grimshaw JR, Halsey M, Sullivan KAM, Moreno-Santillán DD, Crookshanks C, Roberts J, Garcia C. et al. 2023. Insights into mammalian TE diversity through the curation of 248 genome assemblies. Science 380(6643), eabn1430. 10.1126/science.abn1430

61. Packiaraj J, Thakur J. 2024. DNA satellite and chromatin organization at mouse centromeres and pericentromeres. Genome Biol 25:52. 10.1186/s13059-024-03184-z

62. Phifer-Rixey M, Bomhoff M, Nachman MW. 2014. Genome-wide patterns of differentiation among house mouse subspecies inferred from transcriptome sequencing. Genetics 198:283–297.

63. Phifer-Rixey M, Bi K, Ferris KG, Sheehan MJ, Lin D, Mack KL, Keeble SM, Suzuki TA, Good JM, Nachman MW. 2018. The genomic basis of environmental adaptation in house mice. PLoS Genet 14(9): e1007672. 10.1371/journal.pgen.1007672

64. Poplin R, Chang PC, Alexander D, Schwartz S, Colthurst T, Ku A, Newburger D, Dijamco J, Nguyen N, Afshar PT. 2018. A universal SNP and small-indel variant caller using deep neural networks. Nat Biotech 36, 983–987. 10.1038/nbt.4235

65. Quadrana L, Bortolini Silveira A, Mayhew GF, LeBlanc C, Martienssen RA, Jeddeloh JA, Colot V. 2016. The Arabidopsis thaliana mobilome and its impact at the species level. eLife, 5, e15716. 10.7554/eLife.15716

66. Quadrana L, Henderson IR. 2025. The natural history of transposons in plant pangenomes and panepigenomes. Curr Opin Plant Biol 88: 102818. 10.1016/j.pbi.2025.102818

67. Raingeval M, Leduque B, Baduel P, Edera A, Roux F, Colot V, Quadrana L. 2024. Retrotransposon-driven environmental regulation of FLC leads to adaptive response to herbicide. Nat Plants 10(11):1672–1681. 10.1038/s41477-024-01807-8

68. Rech GE, Bogaerts-Márquez M, Barrón MG, Merenciano M, Villanueva-Cañas JL, Horváth V, Fiston-Lavier AS, Luyten I, Venkataram S, Quesneville H. 2019. Stress response, behavior, and development are shaped by transposable element-induced mutations in Drosophila. PLoS Genet 15(2):e1007900 10.1371/journal.pgen.1007900

69. Rech GE, Radío S, Guirao-Rico S, Aguilera L, Horvath V, Green L, Lindstadt H, Jamilloux V, Quesneville H, González J. 2022. Population-scale long-read sequencing uncovers transposable elements associated with gene expression variation and adaptive signatures in Drosophila. Nat Comm 13:1948. 10.1038/s41467-022-29518-8

70. Richardson SR, Gerdes P, Gerhardt DJ, Sanchez-Luque FJ, Bodea GO, Muñoz-Lopez M, Jesuadian JS, Kempen MHC, Carreira PE. et al. 2017. Heritable L1 retrotransposition in the mouse primordial germline and early embryo. Genome Res 27(8):1395–1405. 10.1101/gr.219022.116.

71. Robinson JT, Thorvaldsdóttir H, Winckler W, Guttman M, Lander ES, Getz G, Mesirov JP. 2011. Integrative Genomics Viewer. Nat Biotech 29(1):24–26. 10.1038/nbt.1754

72. Salcedo T, Geraldes A, Nachman MW. 2007. Nucleotide variation in wild and inbred mice. Genetics 177: 2277–2291.

73. Schlenk TA, Begun DJ. 2004. Strong selective sweep associated with a transposable element insertion in *Drosophila simulants*. PNAS 101(6):1626–1631. 10.1073/pnas.0305400101

74. Shahid S, Slotkin RK. 2020. The current revolution in transposable element biology enabled by long reads. Curr Opin Plant Biol 54:49–56. 10.1016/j.pbi.2019.12.012

75. Shumate A, Salzberg S L. 2021. Liftoff: accurate mapping of gene annotations. Bioinformatics 37(12):1639–1643. 10.1093/bioinformatics/btaa1016

76. Sim SB, Corpuz RL, Simmonds TJ, Geib SM. 2022. HiFiAdapterFilt, a memory efficient read processing pipeline. BMC Genomics 23:157. 10.1186/s12864-022-08375-1

77. Simão FA, Waterhouse RM, Ioannidis P, Kriventseva EV, Zdobnov EM. 2015. BUSCO: assessing genome assembly and annotation completeness. Bioinformatics 31(19):3210– 3212. 10.1093/bioinformatics/btv351

78. Smith A, Hubley R. 2015. RepeatMasker. http://www.repatmasker.org

79. Smolka M, Paulin LF, Grochowski CM, Horner DW, Mahmoud M, Behera S, Kalef-Ezra E, Gandhi M, Hong K, Pehlivan D, Scholz SW. 2024. Detection of mosaic and population-level structural variants with Sniffles2. Nat Biotech 42(10):1571–1580. 10.1038/s41587-023-02024-y

80. Storer J, Hubley R, Rosen J, Wheeler TJ, Smit AF. 2021. The Dfam community resource of transposable element families. Mobile DNA, 12, 2. 10.1186/s13100-020-00230-y

81. Sudmant PH, Rausch T, Gardner EJ, Handsaker RE, Abyzov A, Huddleston J, Zhang Y, Ye K, Jun G, Fritz MH. et al. 2015. An integrated map of structural variation in 2,504 human genomes. Nature 7571:75–81. 10.1038/nature15394

82. Sundaram V. Wysocka J. 2020. Transposable elements as a potent source of diverse *cis*-regulatory sequences in mammalian genomes. Philos Trans R Soc Lond B Biol Sci 1795: 20190347. 10.1098/rstb.2019.0347

83. Uliano-Silva M, Ferreira JGRN, Krasheninnikova K; Darwin Tree of Life Consortium; Formenti G, Abueg L, Torrance J, Myers EW, Durbin R, Blaxter M, McCarthy SA. 2023. MitoHiFi: a python pipeline for mitochondrial genome assembly. BMC Bioinformatics 24:288. 10.1186/s12859-023-05385-y

84. Ullastres A, Merenciano M, González J. 2021. Regulatory regions in natural transposable element insertions drive interindividual differences in response to immune challenges in *Drosophila*. Genome Biol 22(265). 10.1186/s13059-021-02471-3

85. Van’t Hof AE, Campagne P, Rigden DJ, Yung CJ, Lingley J, Quail MA, Hall N, Darby AC, Saccheri IJ. 2016. The industrial melanism mutation in British peppered moths is a transposable element. Nature 7605:102–5. doi: 10.1038/nature17951

86. Vargas-Chavez C, Longo Pendy NM, Nsango SE, Aguilera L, Ayala D, González J. 2022. Transposable element variants and their potential adaptive impact in urban populations of the malaria vector *Anopheles coluzzii*. Genome Res 1:189–202. doi: 10.1101/gr.275761.121.

87. Vaser R, Sović I, Nagarajan N, Šikić M. 2017. Fast and accurate de novo genome assembly from long uncorrected reads. Genome Res 27(5):737–746. 10.1101/gr.214270.116

88. Vorontsov IE, Eliseeva IA, Zinkevich A, Nikonov M, Abramov S, Boytsov A, Kamenets V, Kasianova A, Kolmykov S, Yevshin IS, Favorov A, Medvedeva YA, Jolma A, Kolpakov F, Makeev VJ, Kulakovskiy IV. 2024. HOCOMOCO in 2024: a rebuild of the curated collection of binding models for human and mouse transcription factors. Nucleic Acids Res 52(D1):154–163. 10.1093/nar/gkad1077

89. Wicker T, Sabot F, Hua-Van A, Bennetzen JL, Capy P, Chalhoub B, Flavell A, Leroy P, Morgante M, Panaud O. 2007. A unified classification system for eukaryotic transposon elements. Nat Rev Genet 8:973–982. 10.1038/nrg2165

90. Yang H, Wang JR, Didion JP, Buus RJ, Bell TA, Welsh CE, Bonhomme F, Yu AH, Nachman MW, Pialek J, Tucker P. 2011. Subspecific origin and haplotype diversity in the laboratory mouse. Nat Genet 43:648–655.

91. Yang X, Lee WP, Ye K, Lee C. 2019. One reference genome is not enough. Genome biol 20(1):104. 10.1186/s13059-019-1717-0

92. Zhang Y, Tautz D. 2021. Adaptive evolution of transposable elements in natural populations. Genom Biol Evol 13(7): evab123. 10.1093/gbe/evab123

93. Zhou Q, Liu M, Xia X, Gong T, Feng J, Liu W, Liu Y, Zhen B, Wang Y, Ding, C. et al. 2017. A mouse tissue transcription factor atlas. Nat Comm 8: 15089.

